# EEG correlates of auditory rise time processing: A systematic review

**DOI:** 10.64898/2026.03.06.710012

**Authors:** Victoria Manasevich, Daria Kostanian, Anton Rogachev, Olga Sysoeva

## Abstract

Rise time (RT) is considered to be one of the most significant acoustical characteristics of auditory speech stimuli. A substantial amount of data has been accumulated on the neurophysiological mechanisms of RT processing under different conditions and in different groups of people, but these data have not been systematised. This review focuses on studies that have investigated electroencephalographic (EEG) markers of RT sensitivity. The present literature search was conducted according to the PRISMA statement in PubMed, Web of Science and APA PsychInfo databases. The resultant review comprised 37 studies that considered diverse aspects of RT processing. The review describes the main stimulation parameters affecting electrophysiological markers of RT processing reflected in different components of event-related potentials, brainstem responses and cortical rhythmic activity. The main finding of this review is that the rise time prolongation leads to a decrease in the amplitude of the main ERP components and an increase in their latencies. However, the sensitivity of the EEG markers varied with the earliest components tracking the subtle difference (few tens of microseconds), while the later components coding the larger one (up to 500 ms). Nevertheless, the observed effects may vary and depend on some aspects of the experimental paradigm, age of participants and speech-related problems. Future research may benefit by addressing understudied clinical groups and ERP components such as P1 and N2, dominated in children.

## Introduction

The ability to process the acoustical characteristics of auditory speech stimuli is an important predictor of speech development. Disturbances in phonological processing may be one of the factors contributing to the development of different speech disorders – i.e., specific learning difficulty that impacts reading and spelling [1,2]. One of the most important acoustical characteristics of auditory speech stimulus is auditory rise time (or rise time, RT). RT physically represents the rates at which amplitude or frequency modulations in the speech signal increase.

The amplitude envelope of a sound describes the variation of its amplitude over time. In terms of perception, shorter amplitude rise time can create a more abrupt and percussive sound, while a longer RT can produce a smoother and more gradual onset, often perceived as softer or more mellow [3]. Very short RTs (<10ms), for example, produce spectral splatter, which is often perceived as a click. Formant RT refers to the duration it takes for the formant frequencies to reach their peak amplitudes after the onset of a vowel sound. The RT of formants can significantly affect speech perception. Formant RT plays a key role in distinguishing between different speech sounds, especially consonants that differ in the way or place of articulation. For example, it helps listeners distinguish between pairs of sounds such as /ba/ and /wa/, where changes in formant frequencies occur at different rates [4]

Psychophysical studies of RT sensitivity provide a framework for assessing the contribution of phonological processing to speech and language development. The difference threshold (i.e., the minimal changes in RT at which stimuli are perceived as different) is used to estimate a measure of RT sensitivity. Individual differences in the ability to discriminate RT may affect the accuracy of linguistic representations, which may be related to individual variation in speech performance. Children with dyslexia have demonstrated less sensitivity to amplitude RT of auditory stimuli than their control group peers [1]. Also, sensitivity to RT in infancy may act as a predictor of vocabulary development in older age [5].

At the neurophysiological level, sensitivity to RT can be reflected in the event-related potentials (ERP) components changes. ERP is a commonly used technique for studying brain responses to stimuli registered with electroencephalogram (EEG). Different ERP components elicited at specific time intervals reflect stages of neurophysiological processing of the stimulus. Both amplitude and formant RTs are crucial for understanding how we perceive sounds and speech. The scientific community has made efforts to generalize some aspects of existing knowledge on RT, yet currently, there is no review that encompasses the mechanisms of RT perception and their reflection in the brain, including both brainstem reactions to short RTs and later cortical responses. А large amount of data has been accumulated on the neurophysiological effects of RT in different groups of people using different methods (ERP, analysis of rhythmic brain activity etc.), but these data are to some degree inconsistent. This inconsistency and the potential role of neurophysiological processing of RT as a possible predictor and marker of speech disorders, creates a need to systematize the empirical data presented in the literature. The aim of this systematic review is to consider evidence of EEG characteristics as markers of sensitivity to RT changes. To provide the most wide-ranging view of the available evidence, this review includes studies considering the neurophysiologic correlates of changes in RT in typically developing children and adults, as well as in groups with different speech disorders, using various approaches to analyze EEG data.

## Methods

This review followed the methodological framework for systematic reviews. To ensure transparent and complete reporting, we adhered to the PRISMA 2020 guidelines. The PRISMA 2020 checklist has been completed and is available as supplementary material, and the PRISMA flow diagram (Figure 1) illustrates the study selection process. While a formal review protocol was not previously registered or published, the search strategy, inclusion criteria, and data extraction forms were developed a priori and piloted by the research team to minimize bias.

**Fig. 1.**
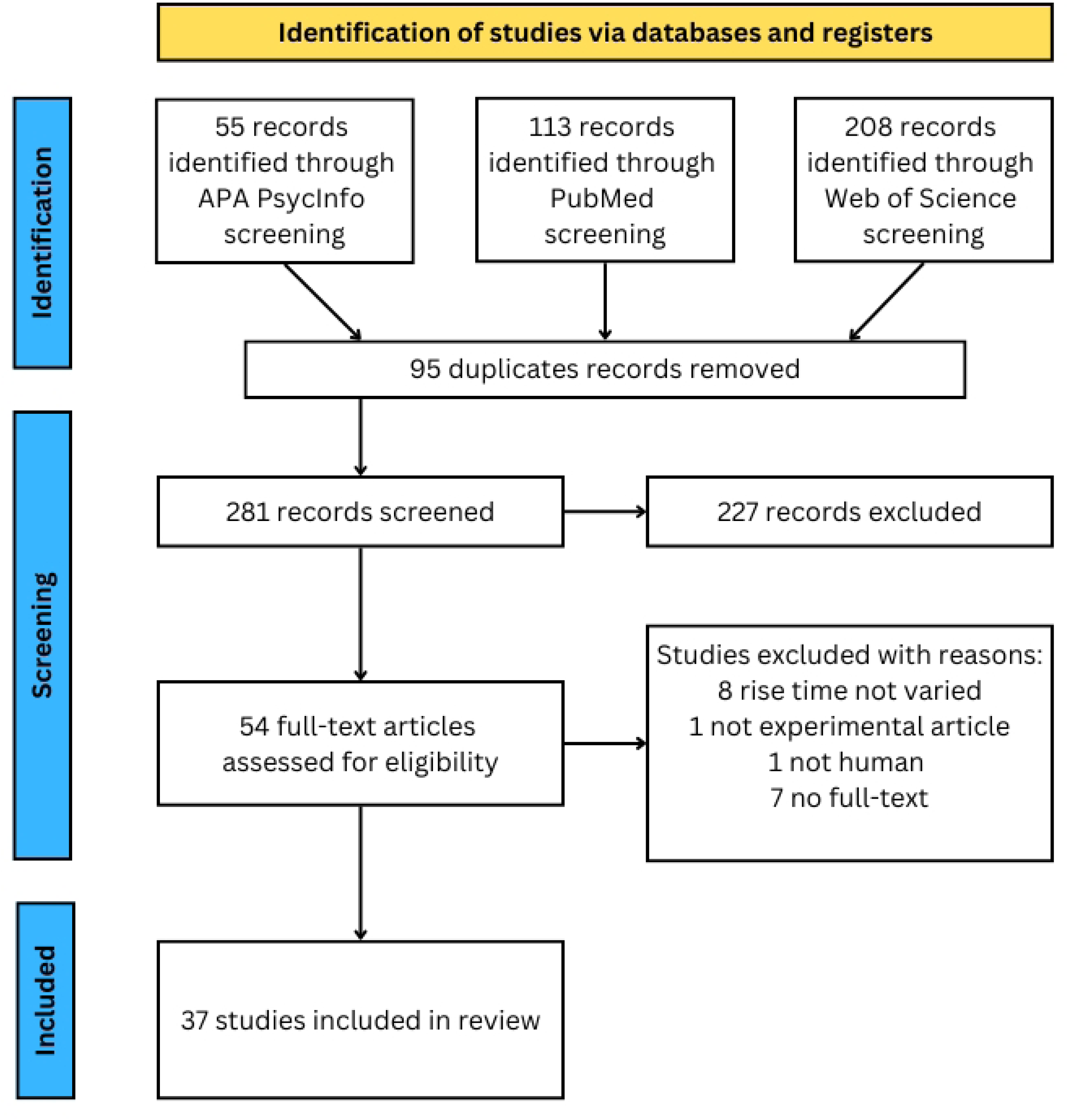
A PRISMA flow-chart representing a flow of information through the different phases of a systematic review.

### Search strategy

We conducted a literature search of the PubMed, Web of Science and APA PsycInfo databases. The search queries used was “(“Rise time”[All Fields] AND (“evoked potentials”[All Fields] OR “erp”[All Fields] OR “evoked response”[All Fields] OR “aep”[All Fields] OR “auditory evoked response”[All Fields] OR “eeg”[All Fields] OR “electrophysiological”[All Fields])) AND (“humans”[Terms] AND “english”[Language]))”. Identified articles were published prior to January 25, 2024. On February 15, 2026, an updated search was performed using the same search strategies and databases as the original search. The search was limited to the period from January 24, 2024, to February 15, 2026, using filters based on entry date. The updated search did not identify any new studies that met the inclusion criteria.

### Eligibility criteria

Only empirical human studies published in the English language in peer-reviewed journals were included in the review. Articles were retained if they measured any ERPs or other EEG data in response to auditory stimuli with varied amplitude/formant RT. Studies were excluded if they contained only one RT condition with no variation.

All titles and abstracts identified in the search were double blind screened by two independent experts using SRDR plus platform (https://srdrplus.ahrq.gov) to exclude obviously irrelevant articles. Any disagreements were resolved through discussion with the involvement of a third expert. The records that passed the initial screening were evaluated using their full-text versions. As in abstract screening, two independent decisions were made for each of the screened items. In case of disagreement, the final decision was made after discussion with the involvement of a third expert. The remaining articles were included in the review.

### Data extraction

The following data were extracted onto a custom sheet, which included information about sample (participant groups, sample size, mean age or age range), experimental paradigm (oddball, block design or other), stimuli information (stimulus type, RT conditions, technical parameters of presentation), additional assessments used in the study (e.g. reading skills assessment or additional deviant condition in oddball study) and results (reported as EEG parameters changes with RT increasing, or observed MMN in response to RT deviants). The extracted data is provided in Table 1.

**Table 1.**
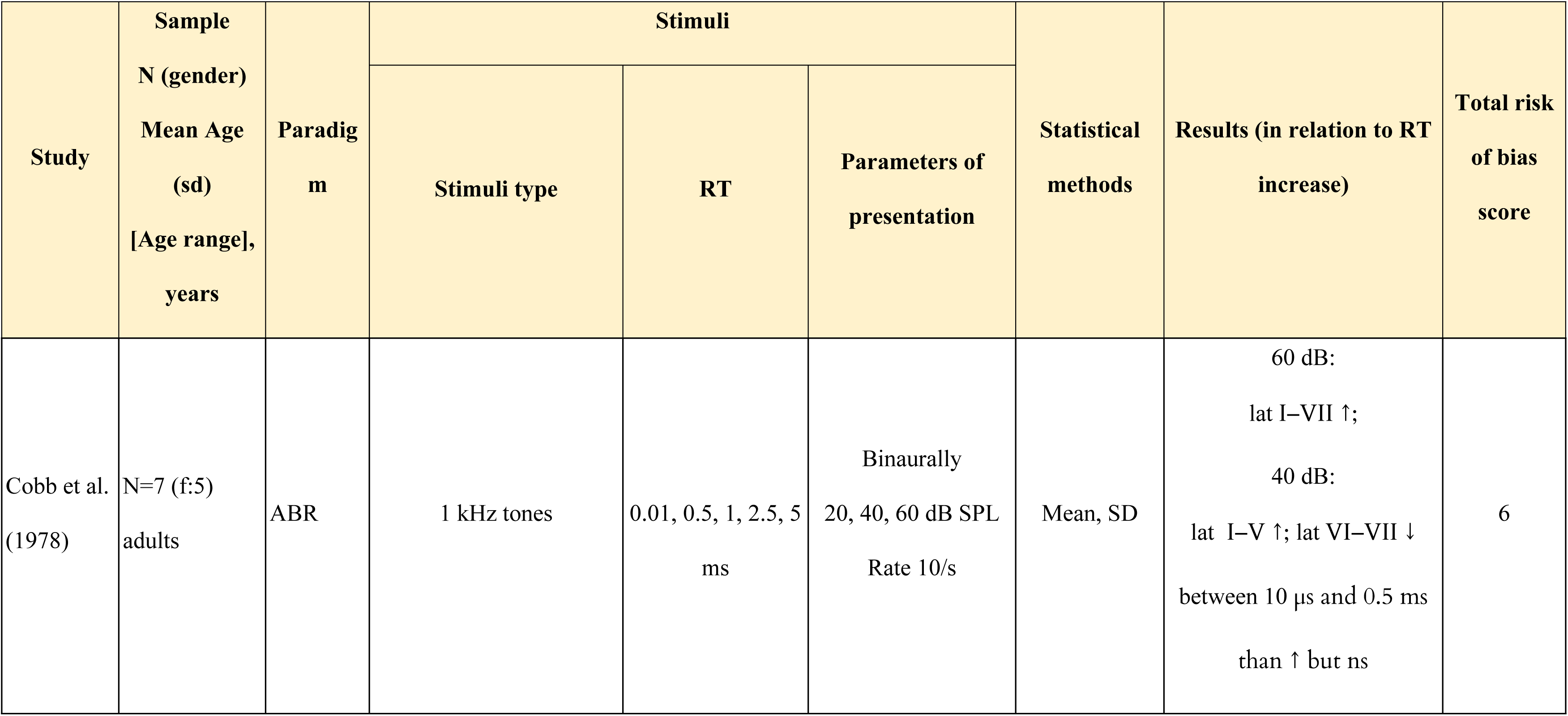
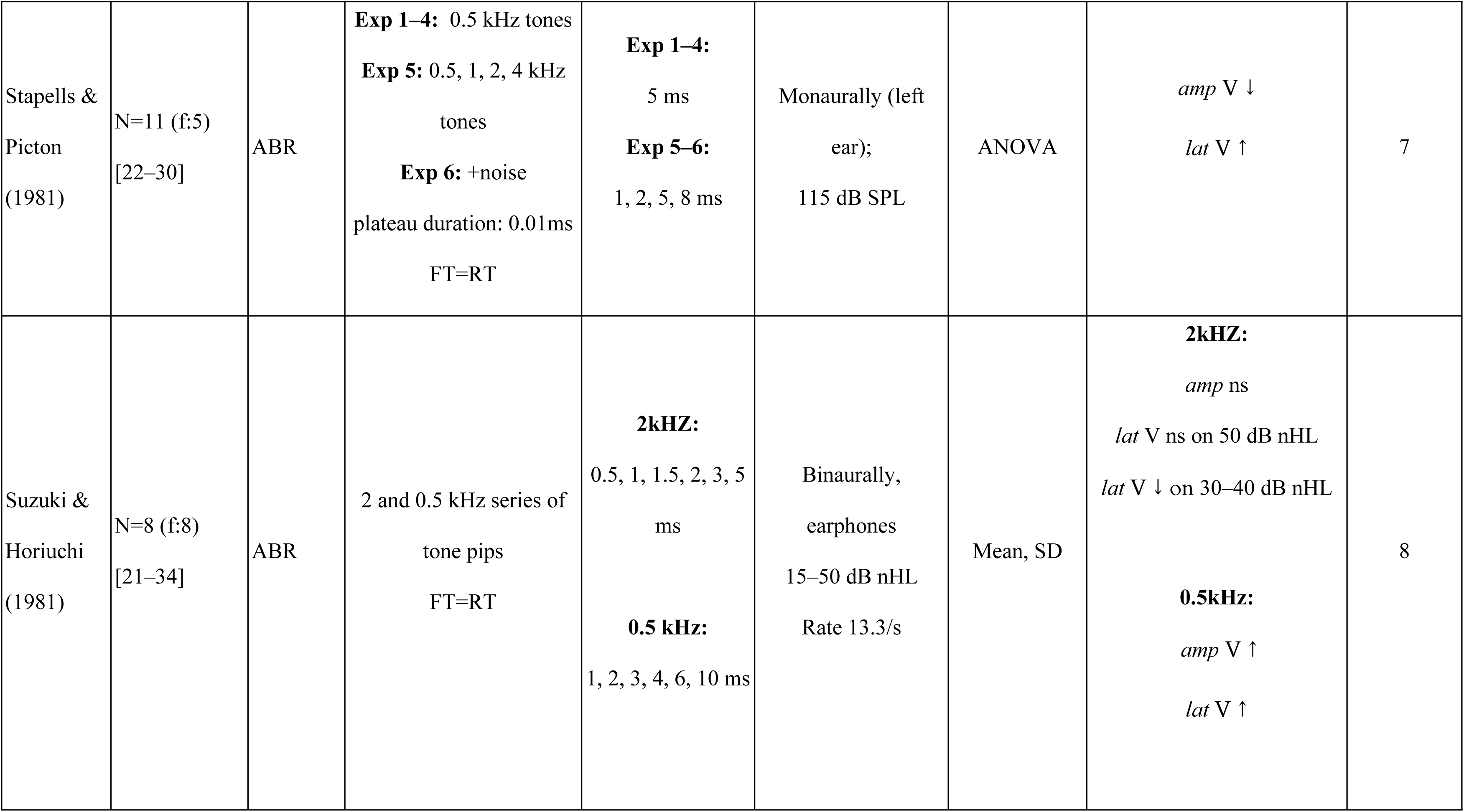
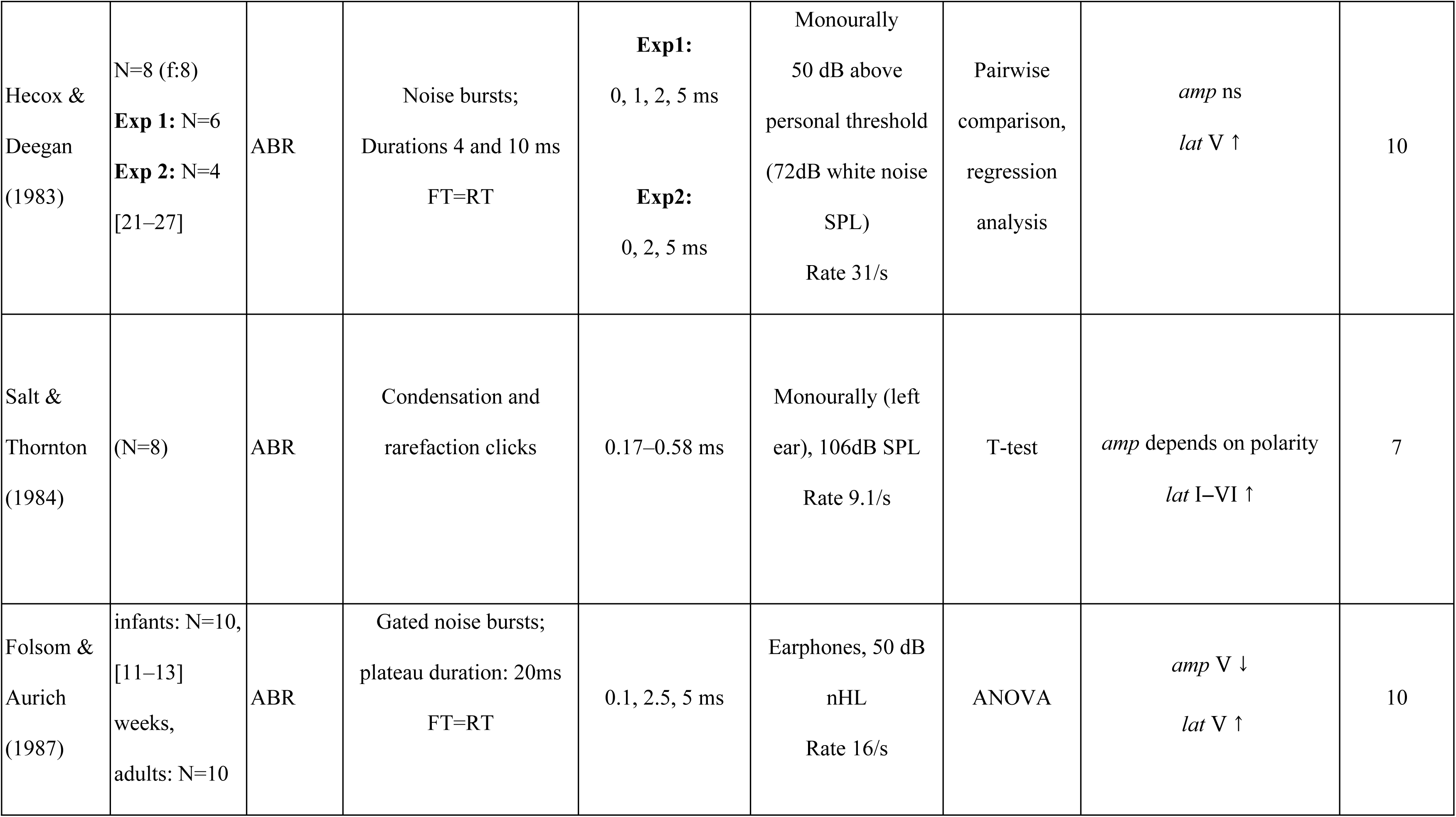
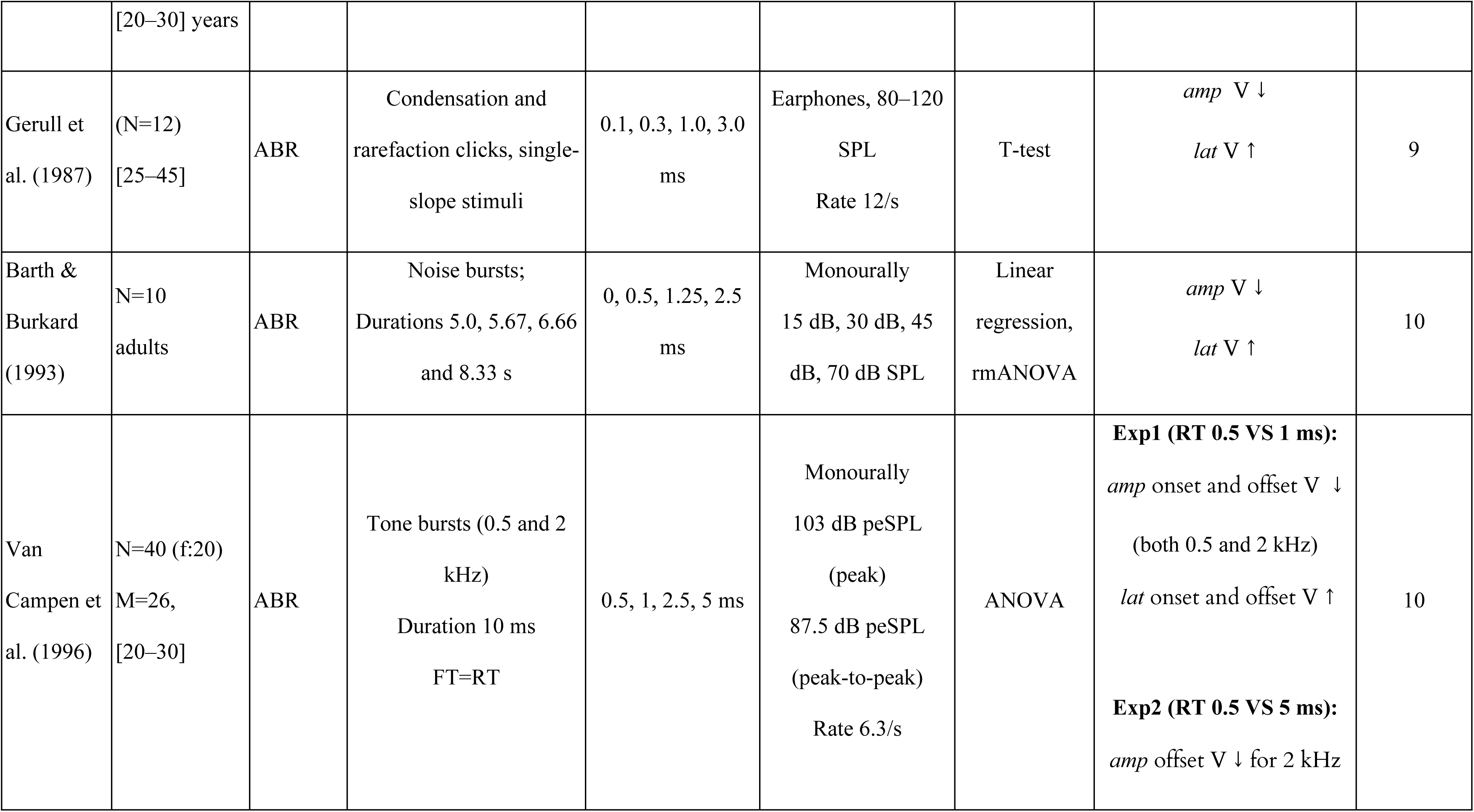
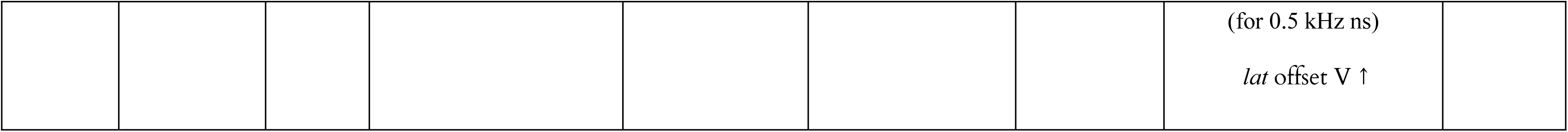
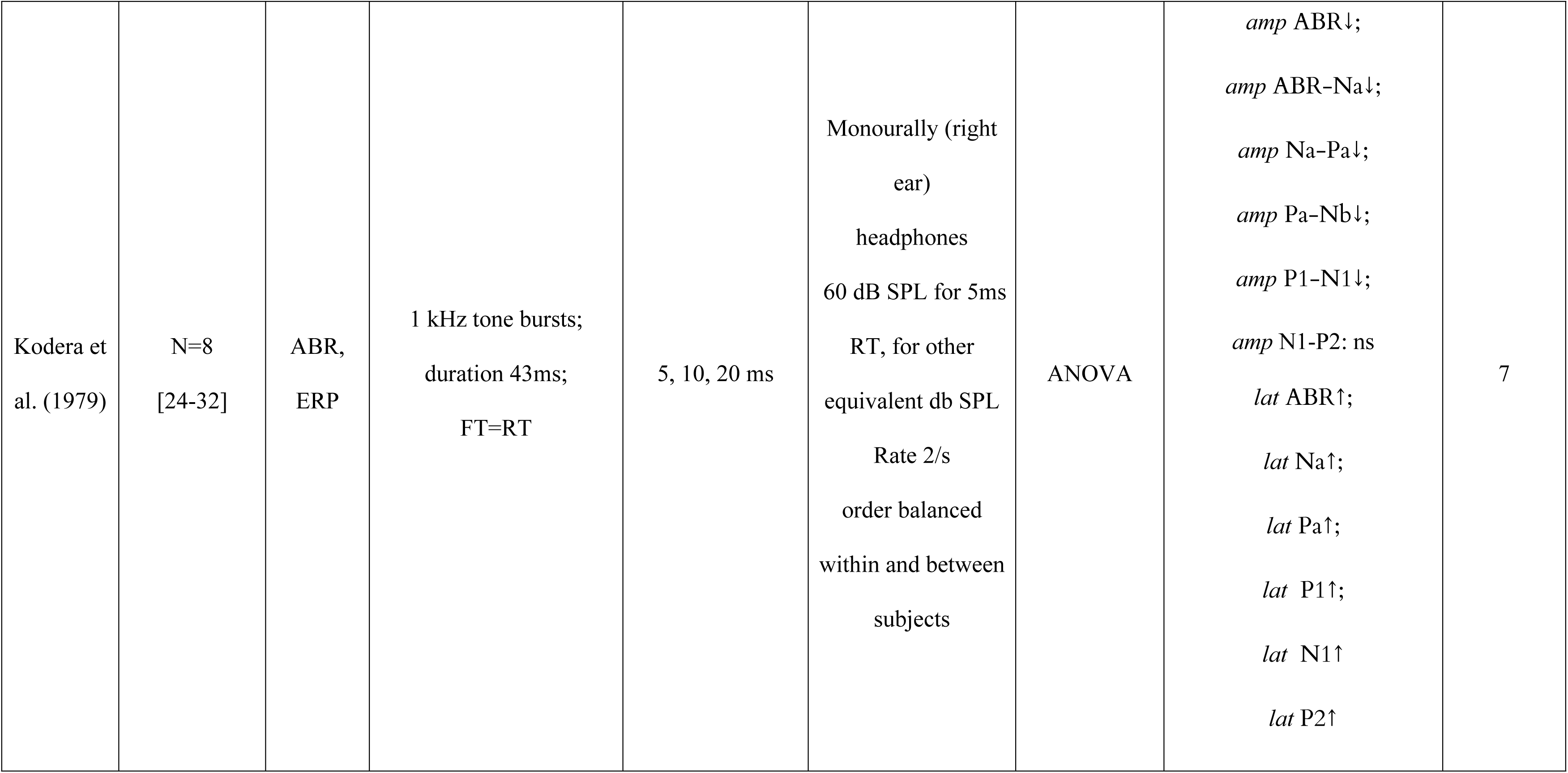
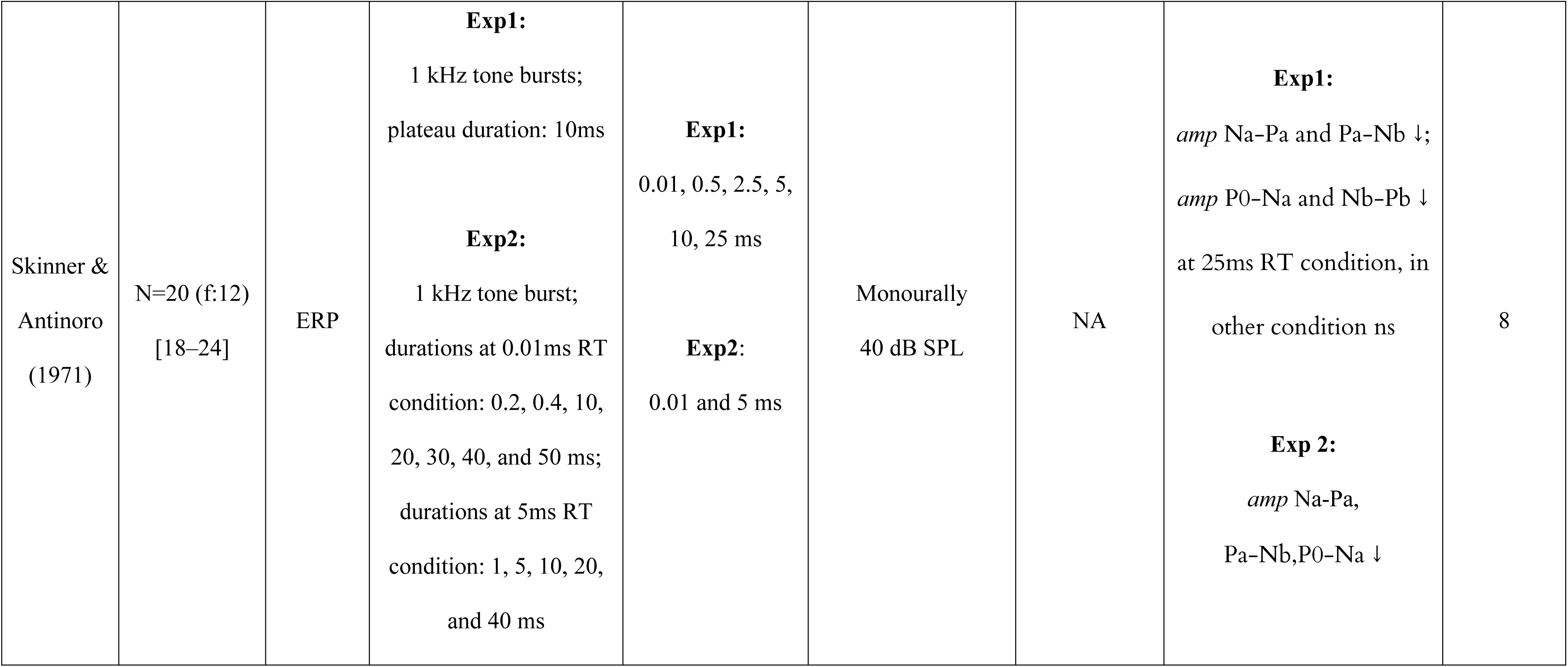
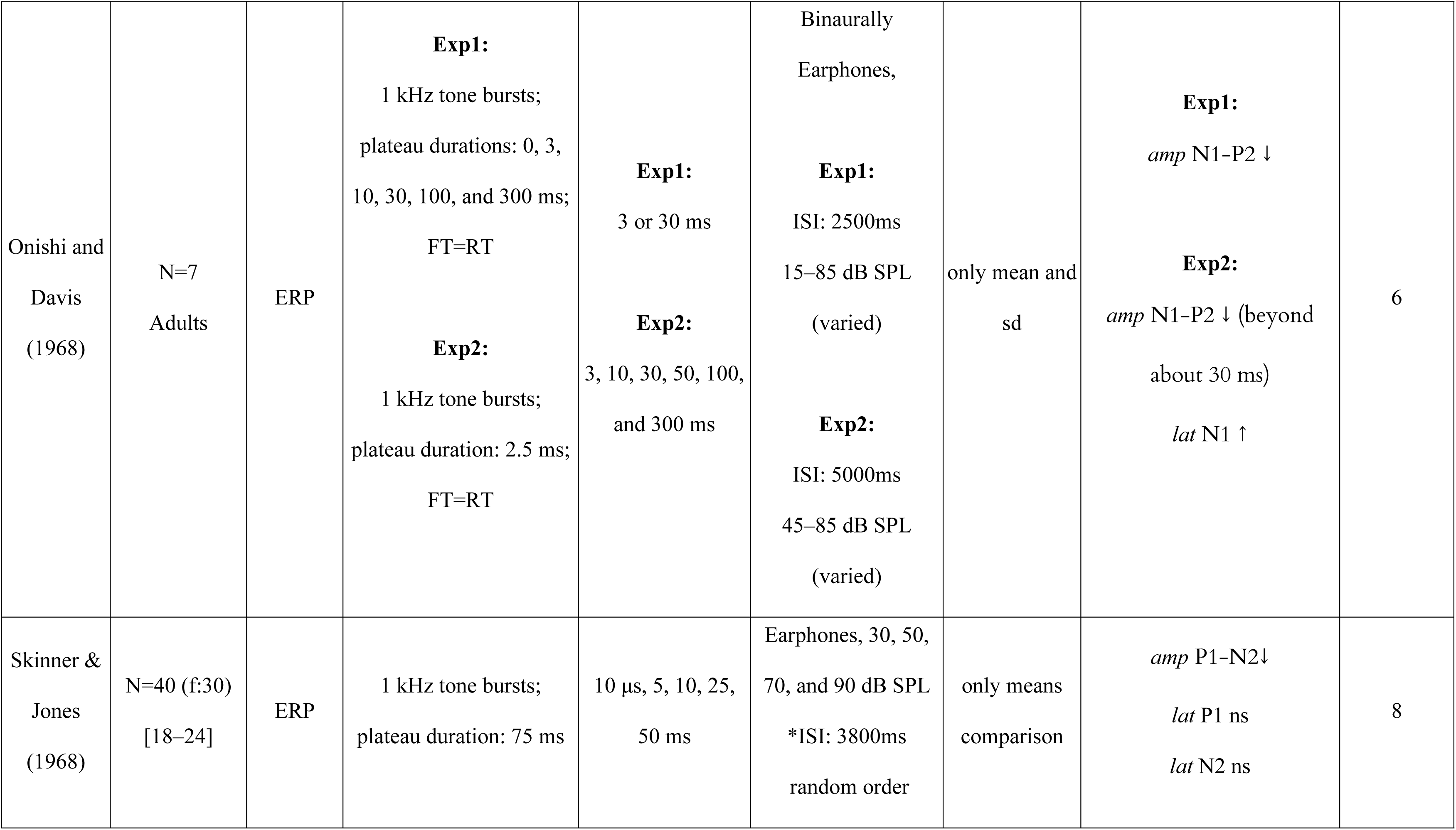
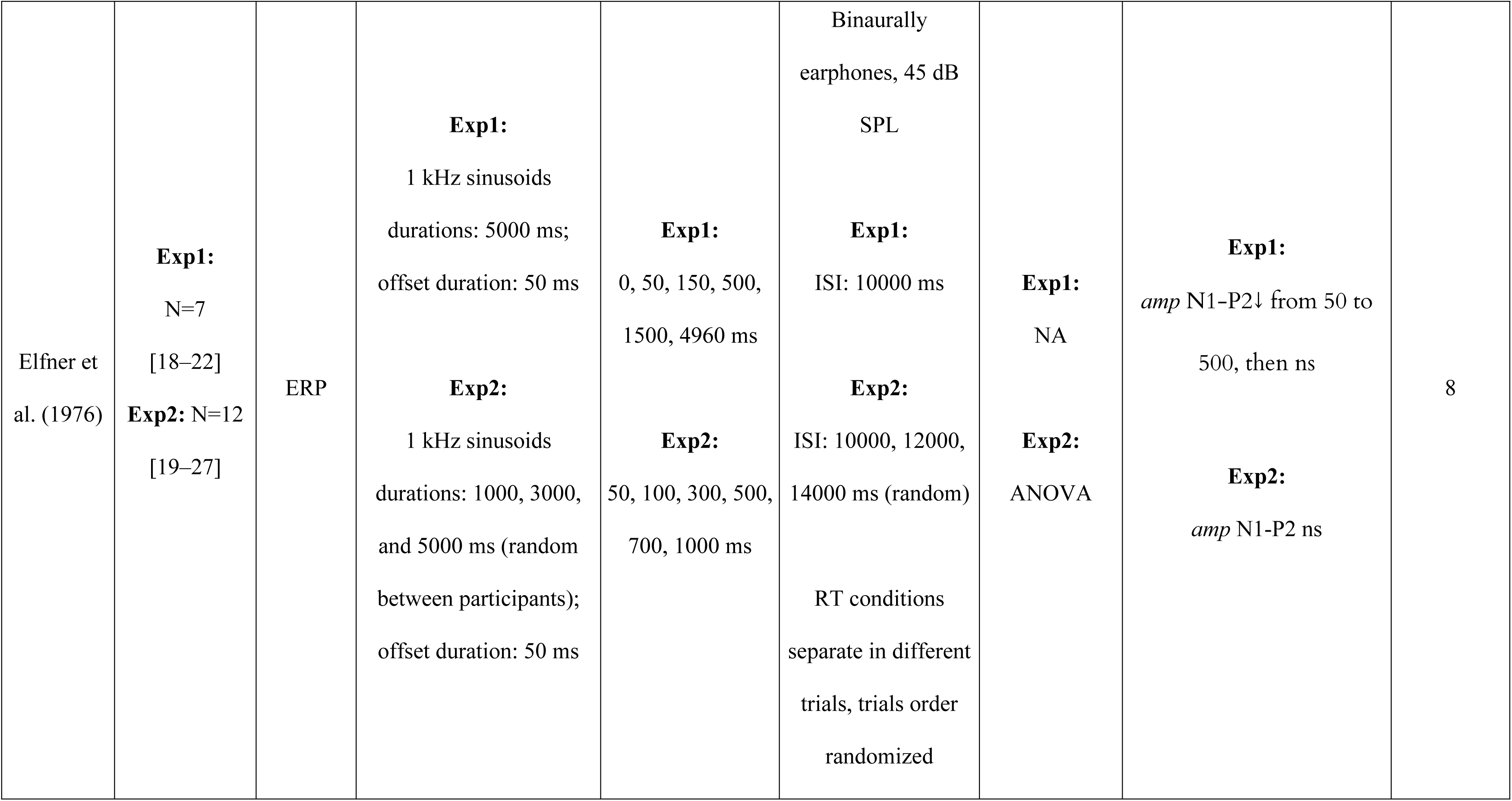
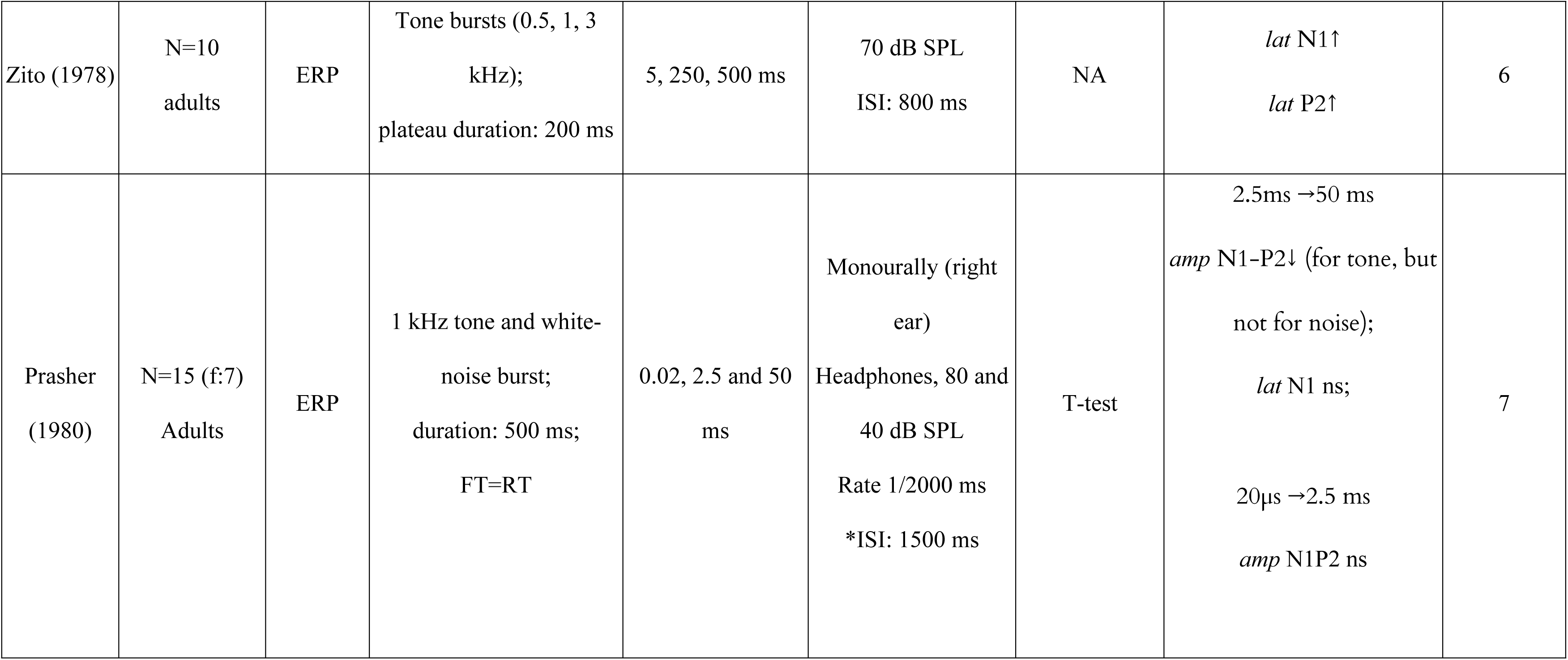
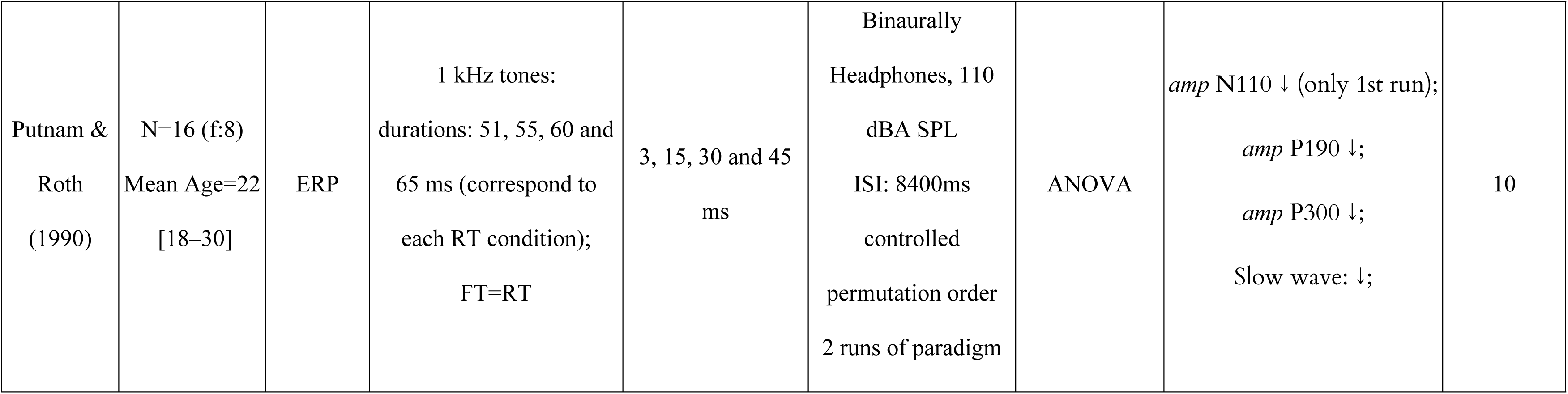
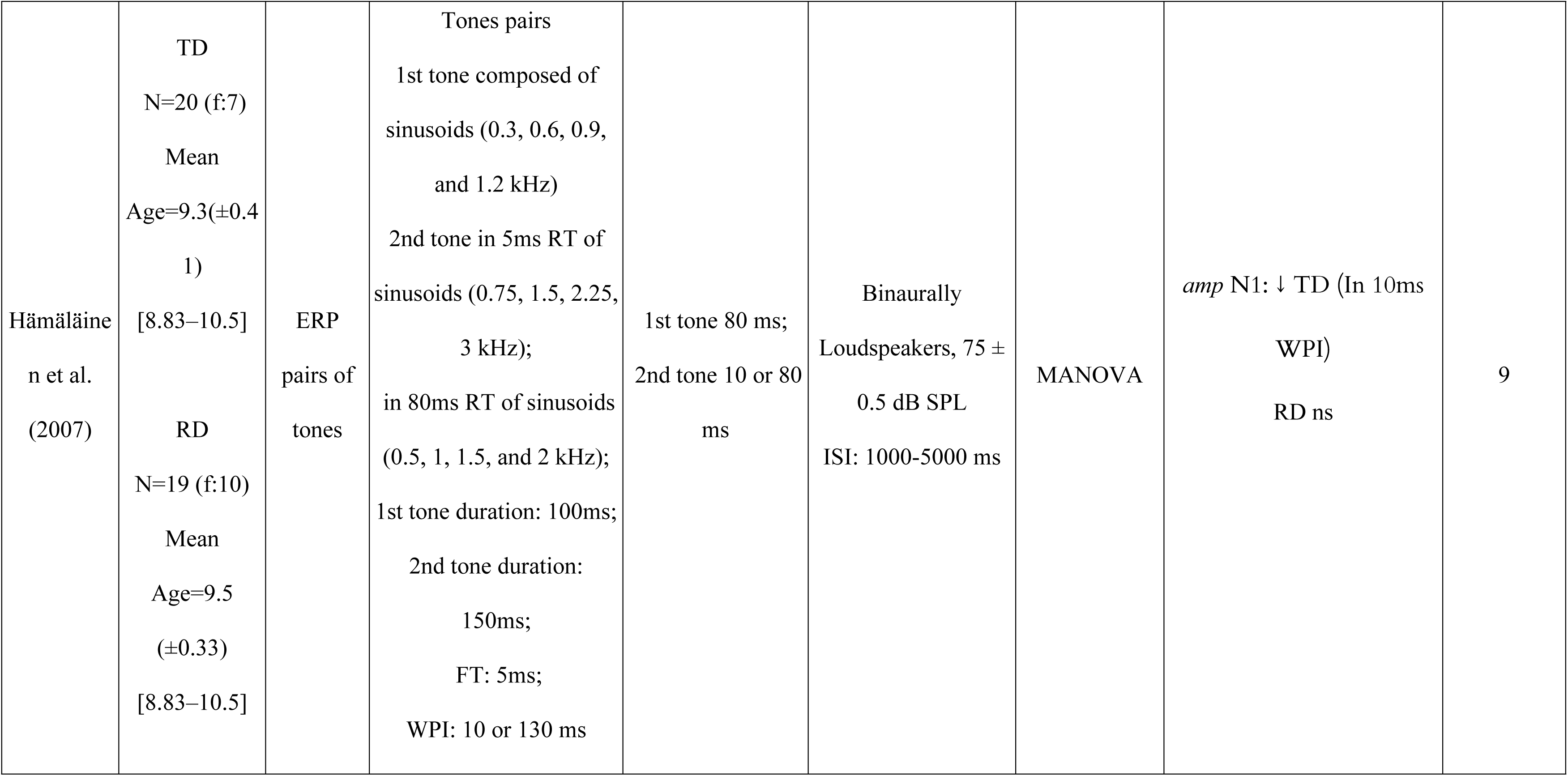
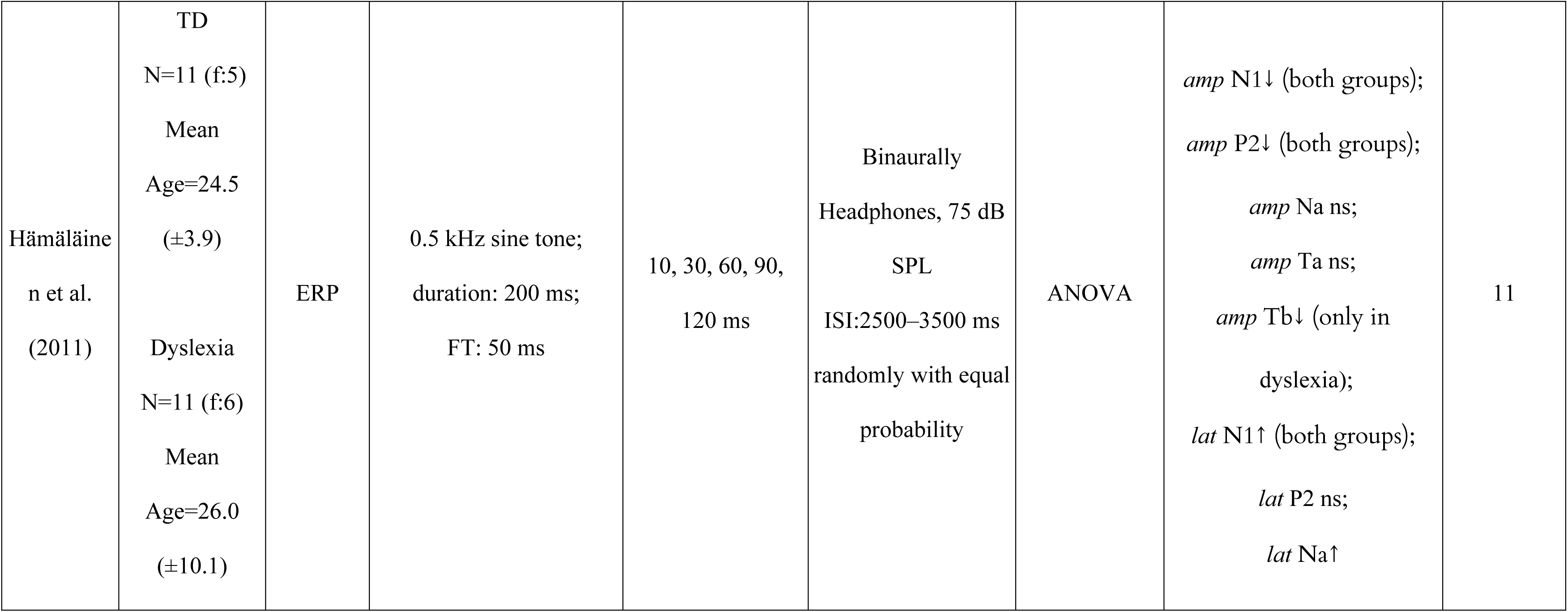
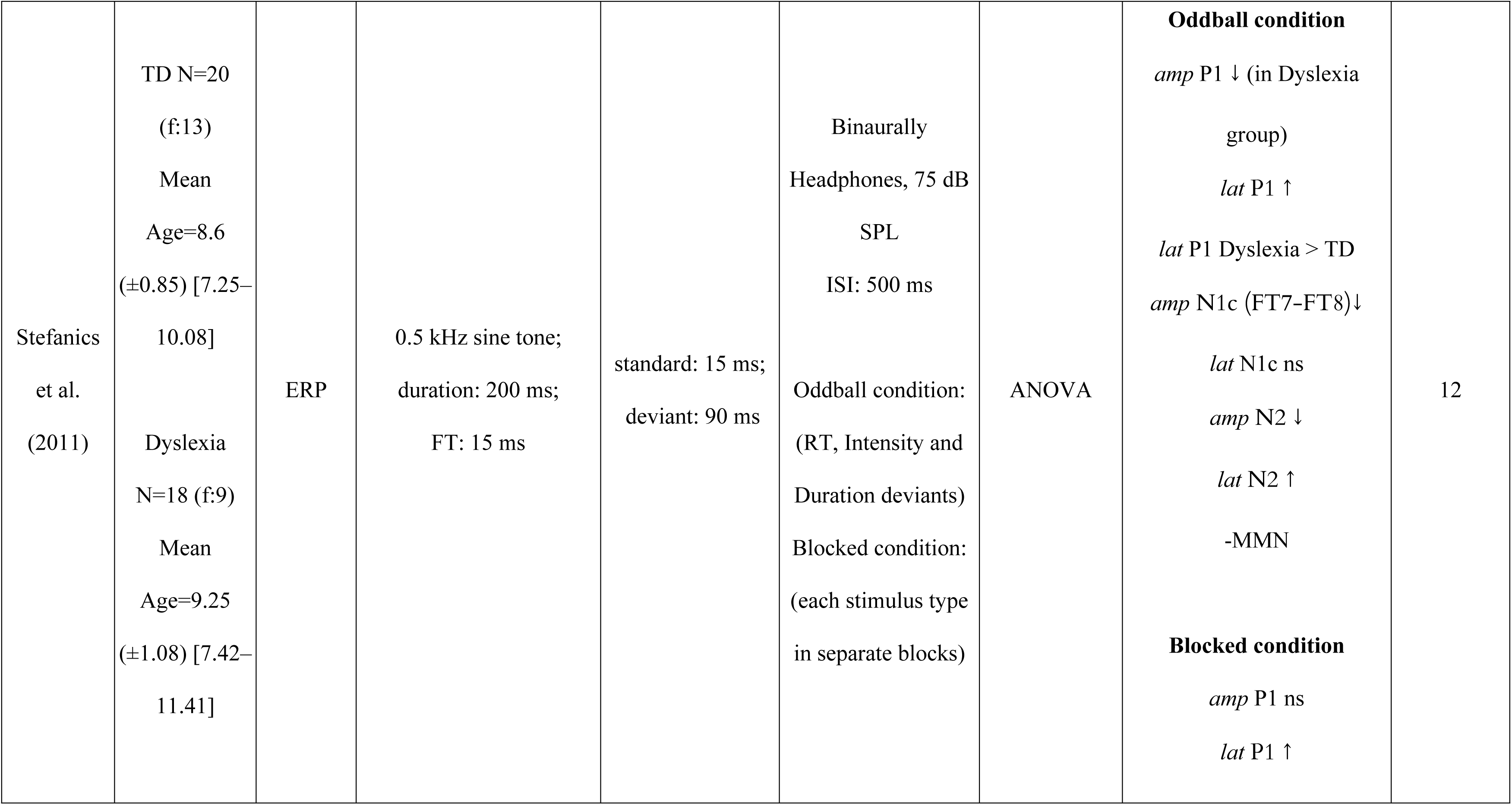
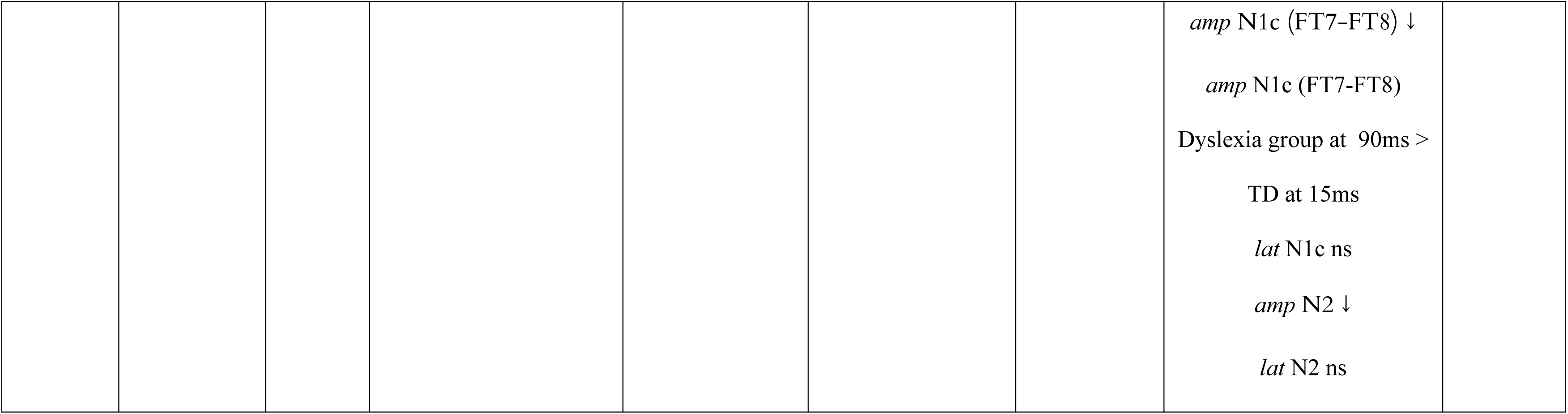
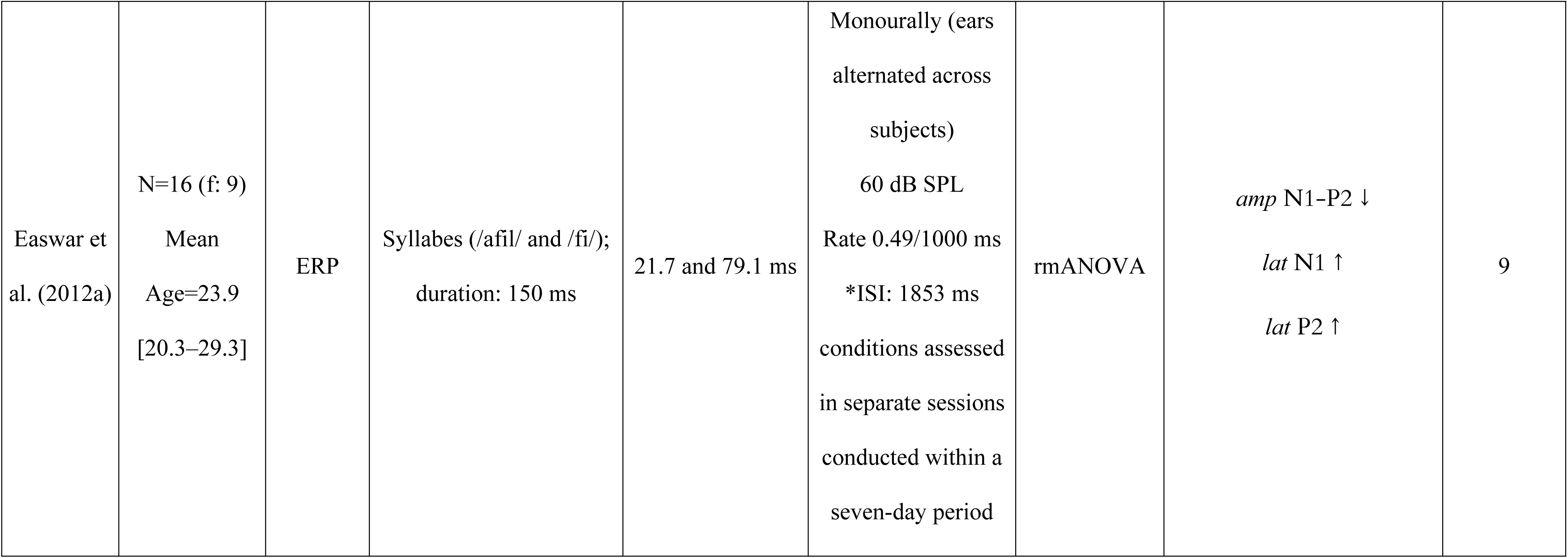
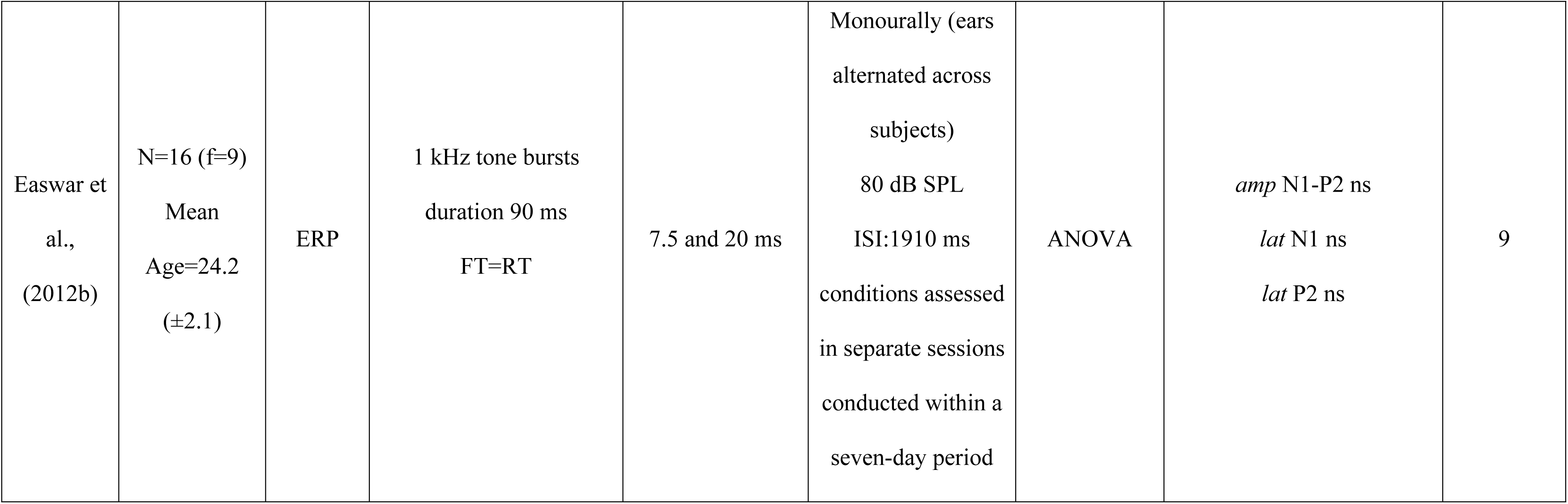
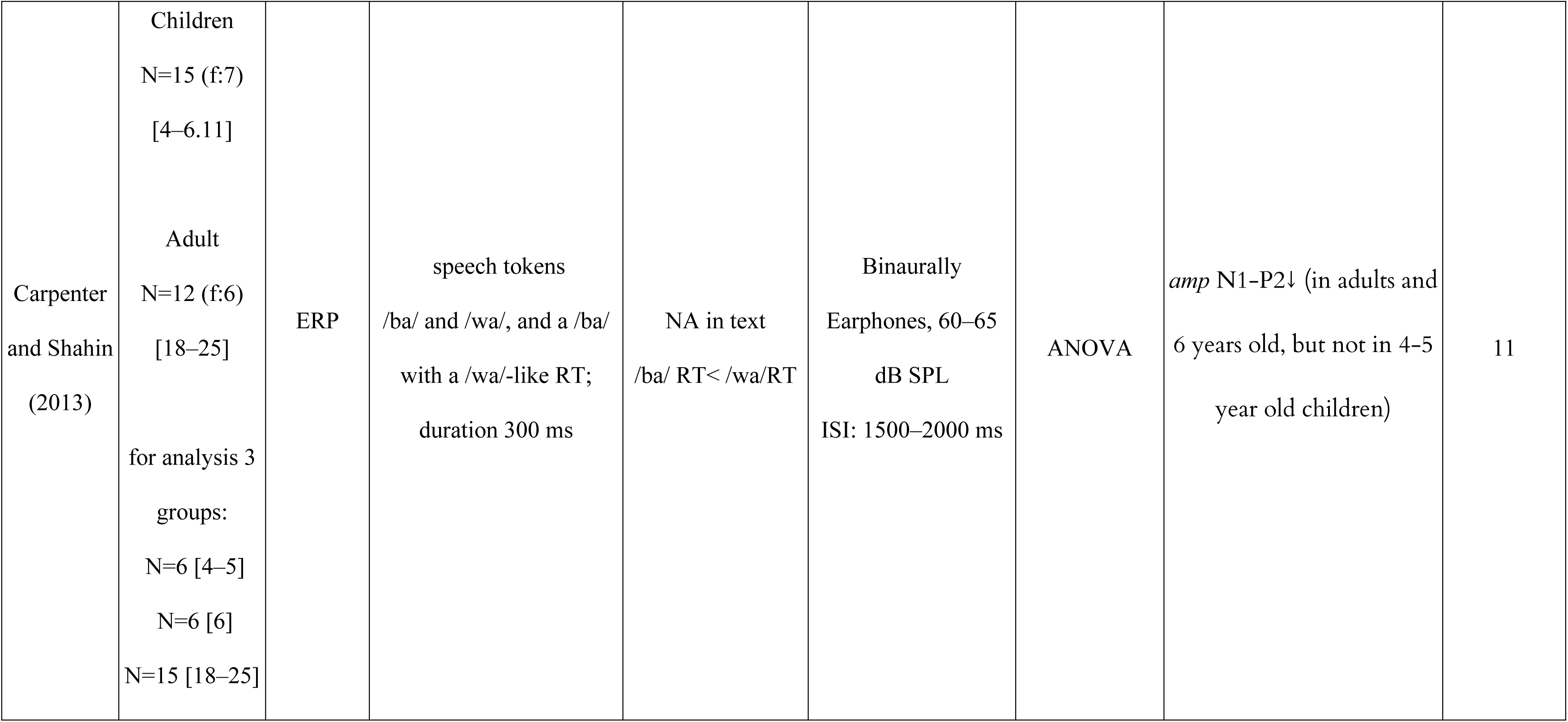
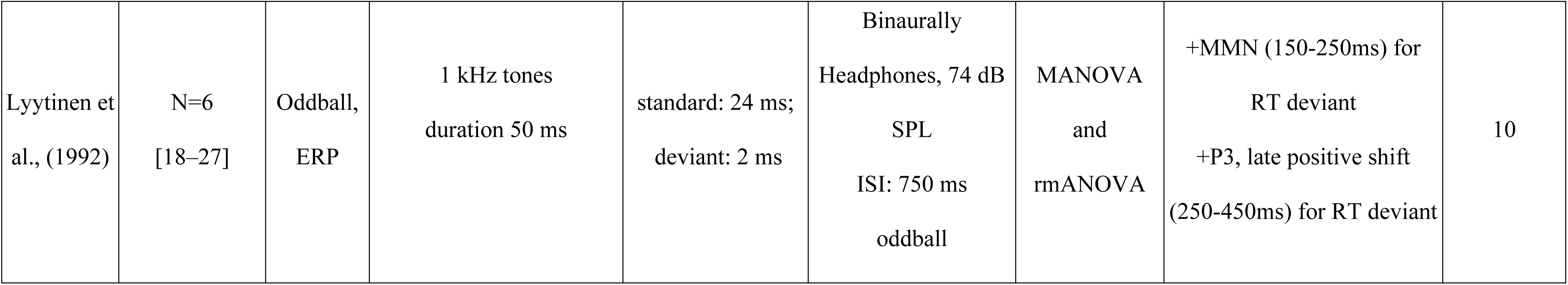
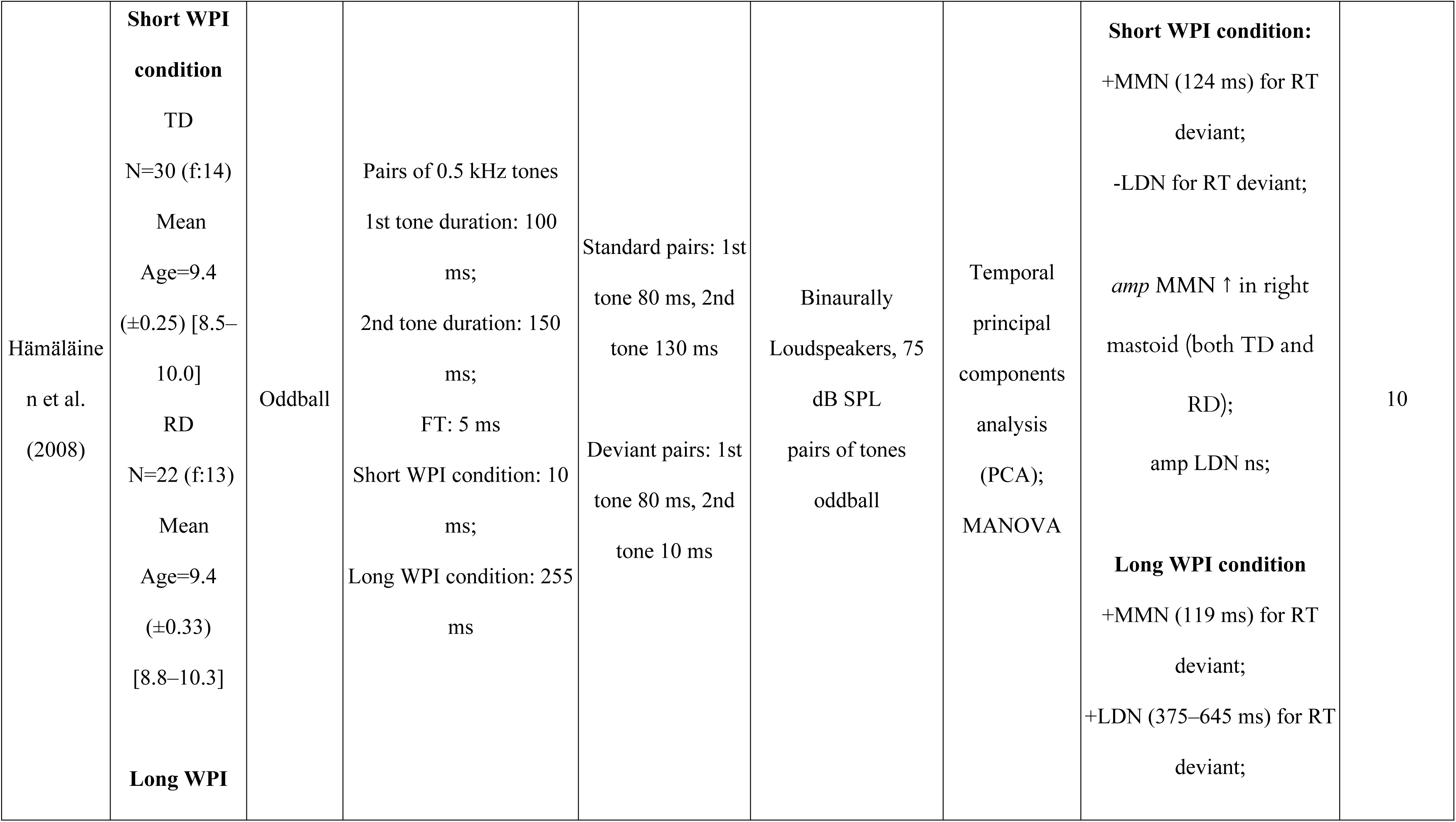
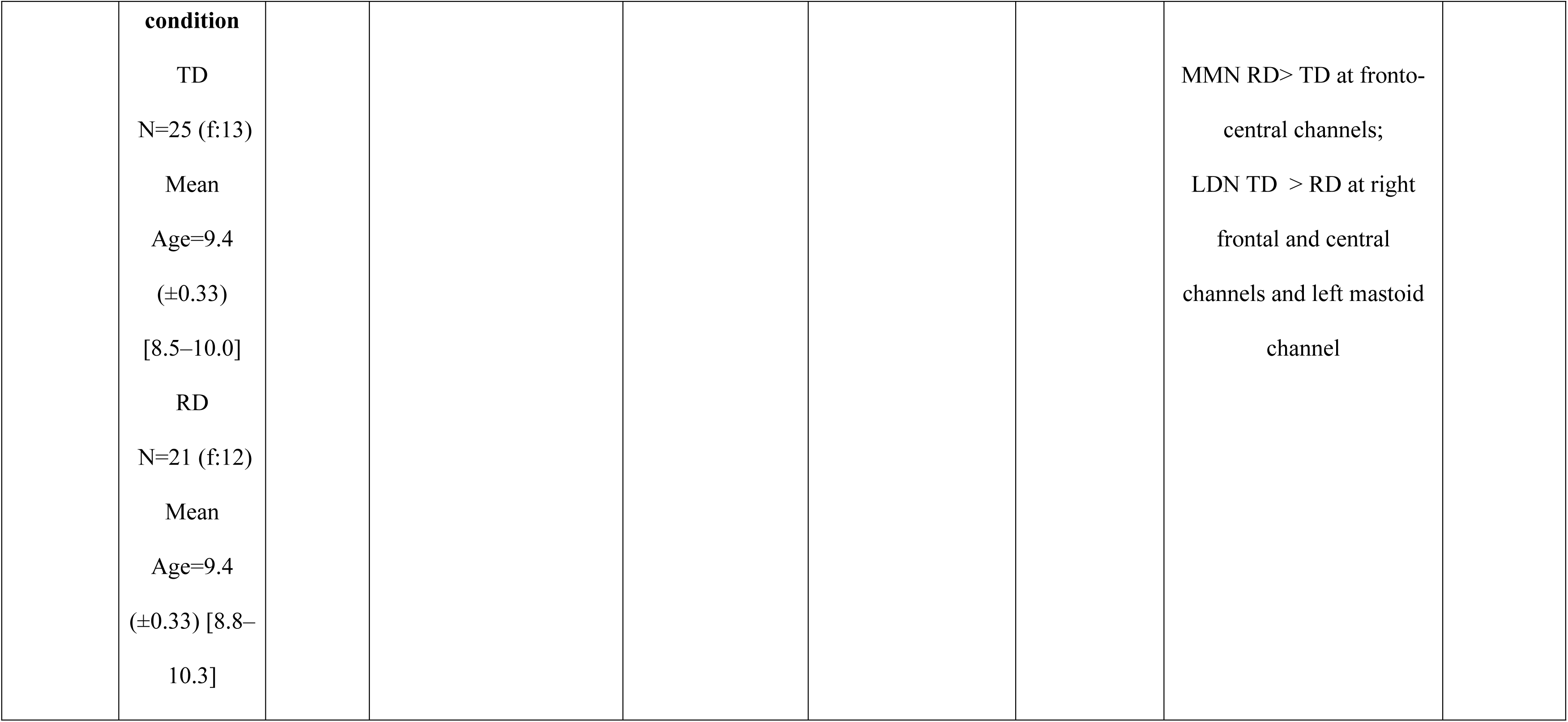
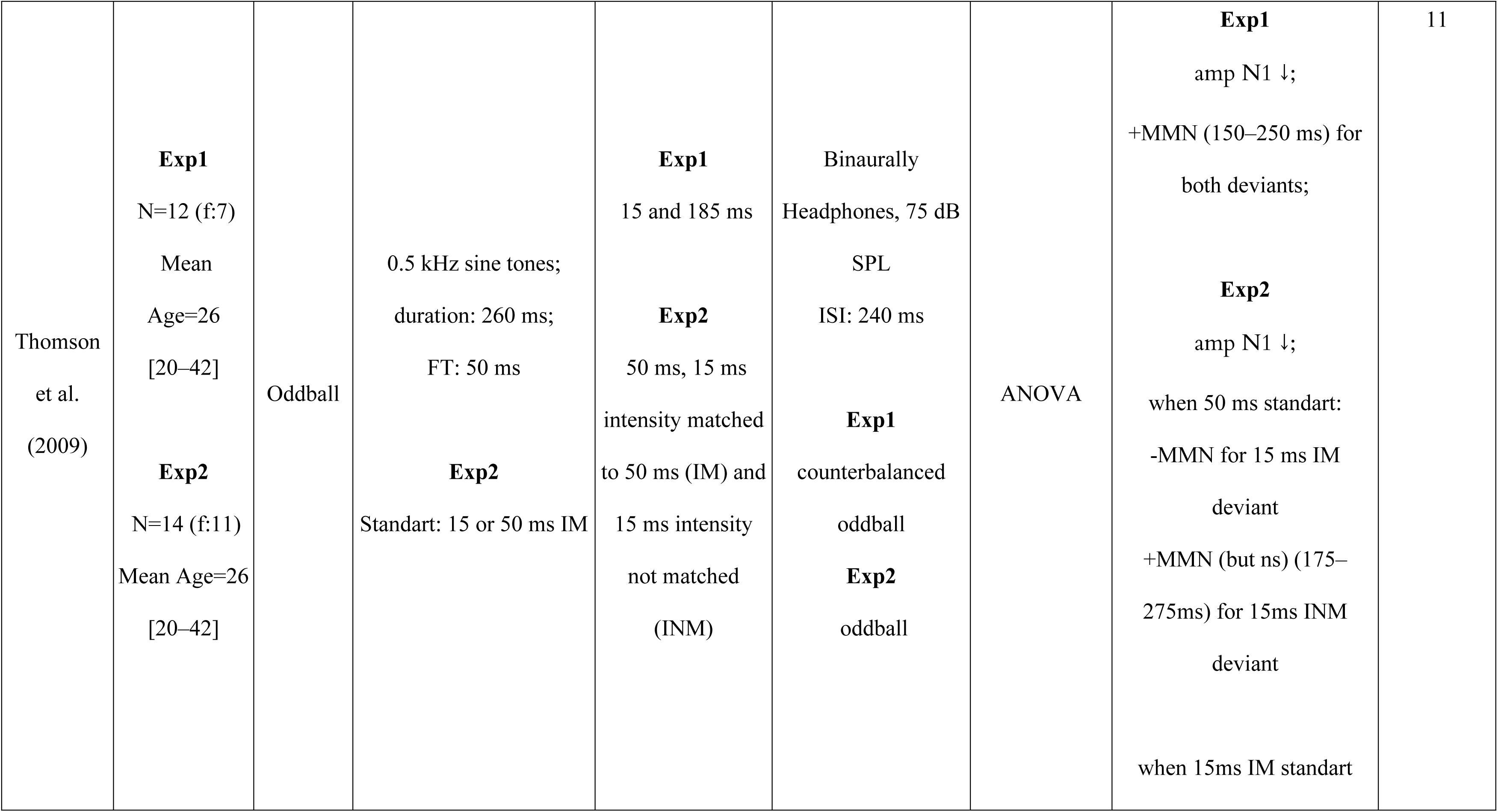
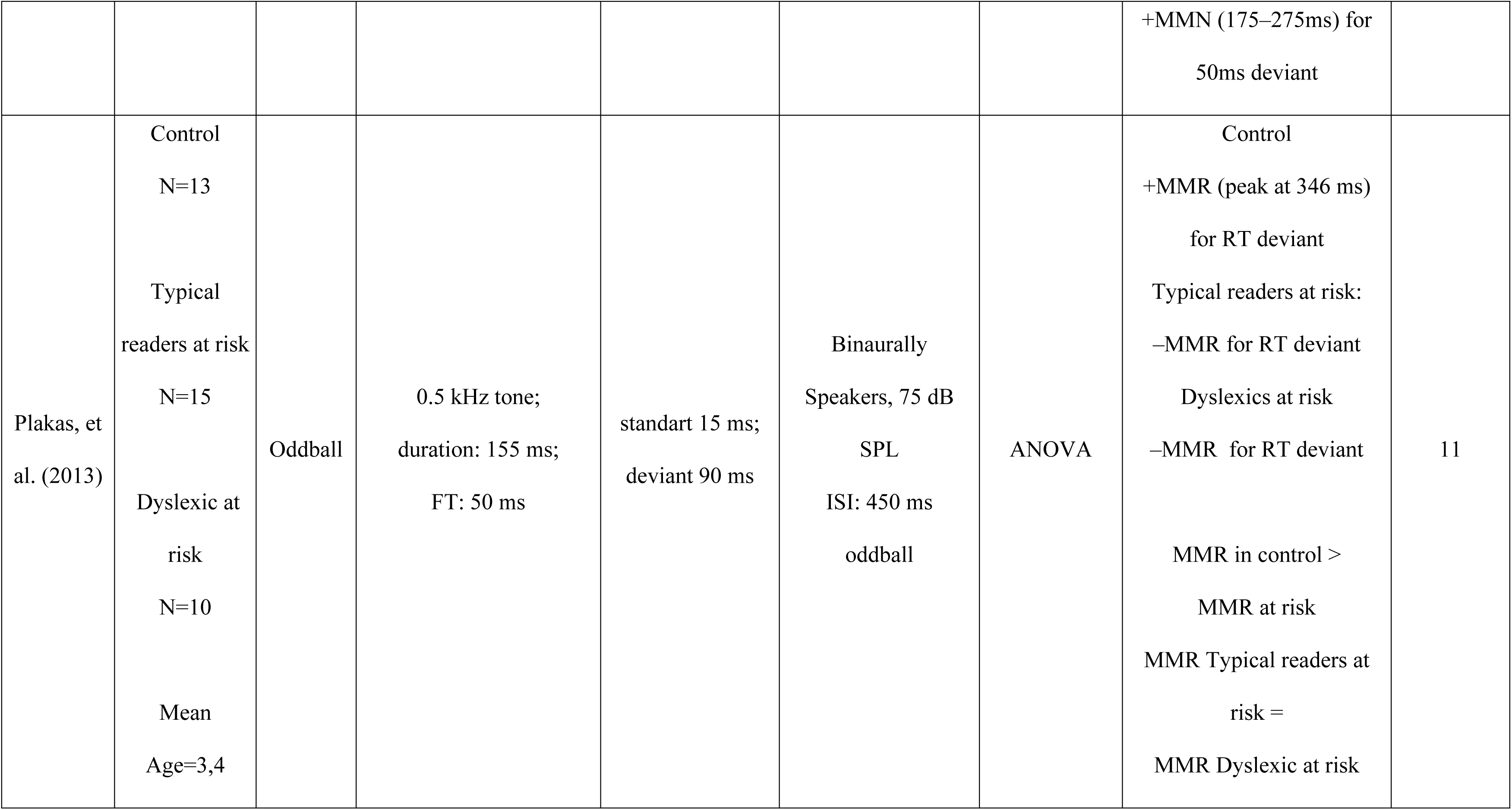
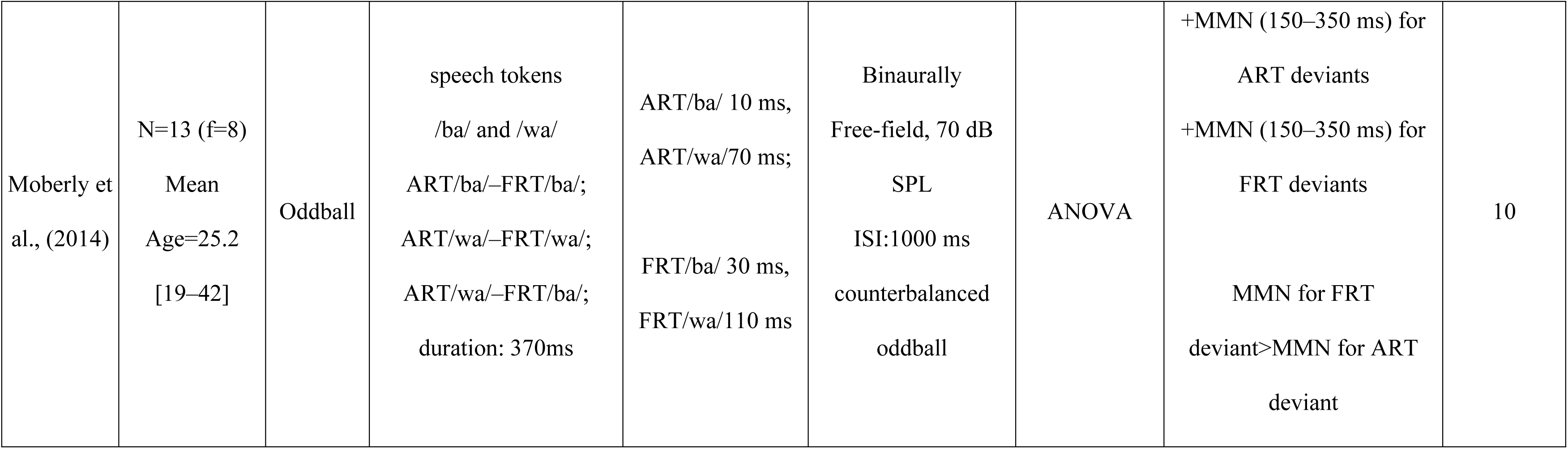
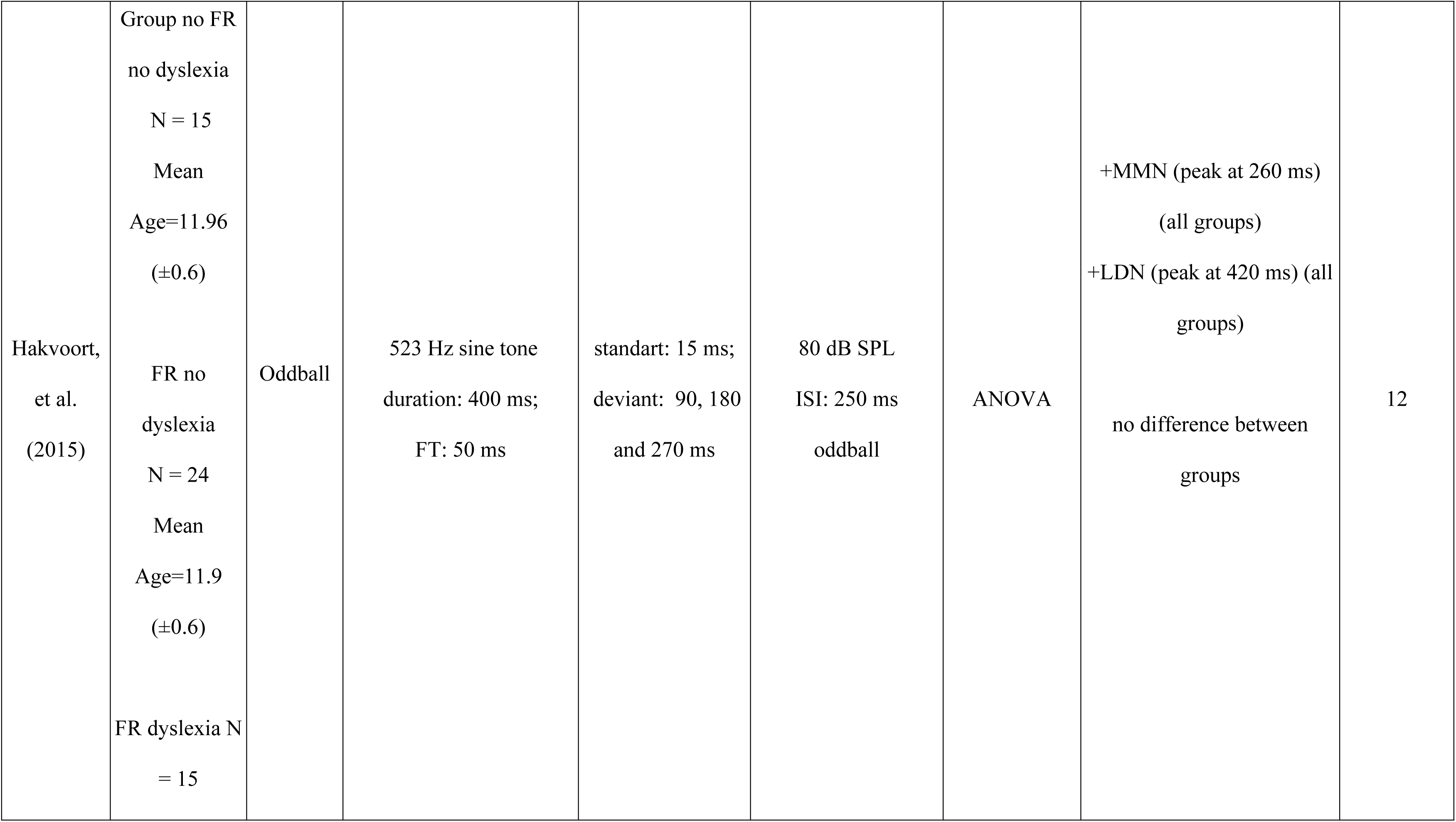
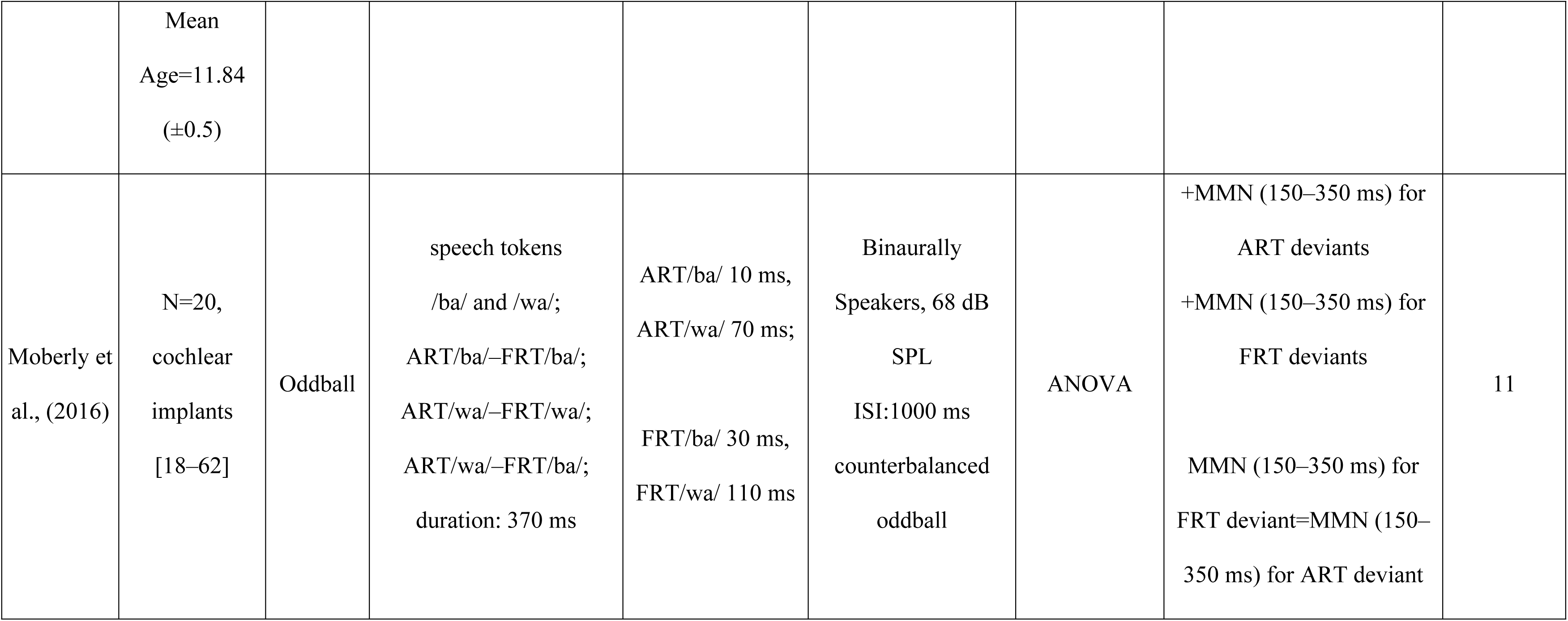
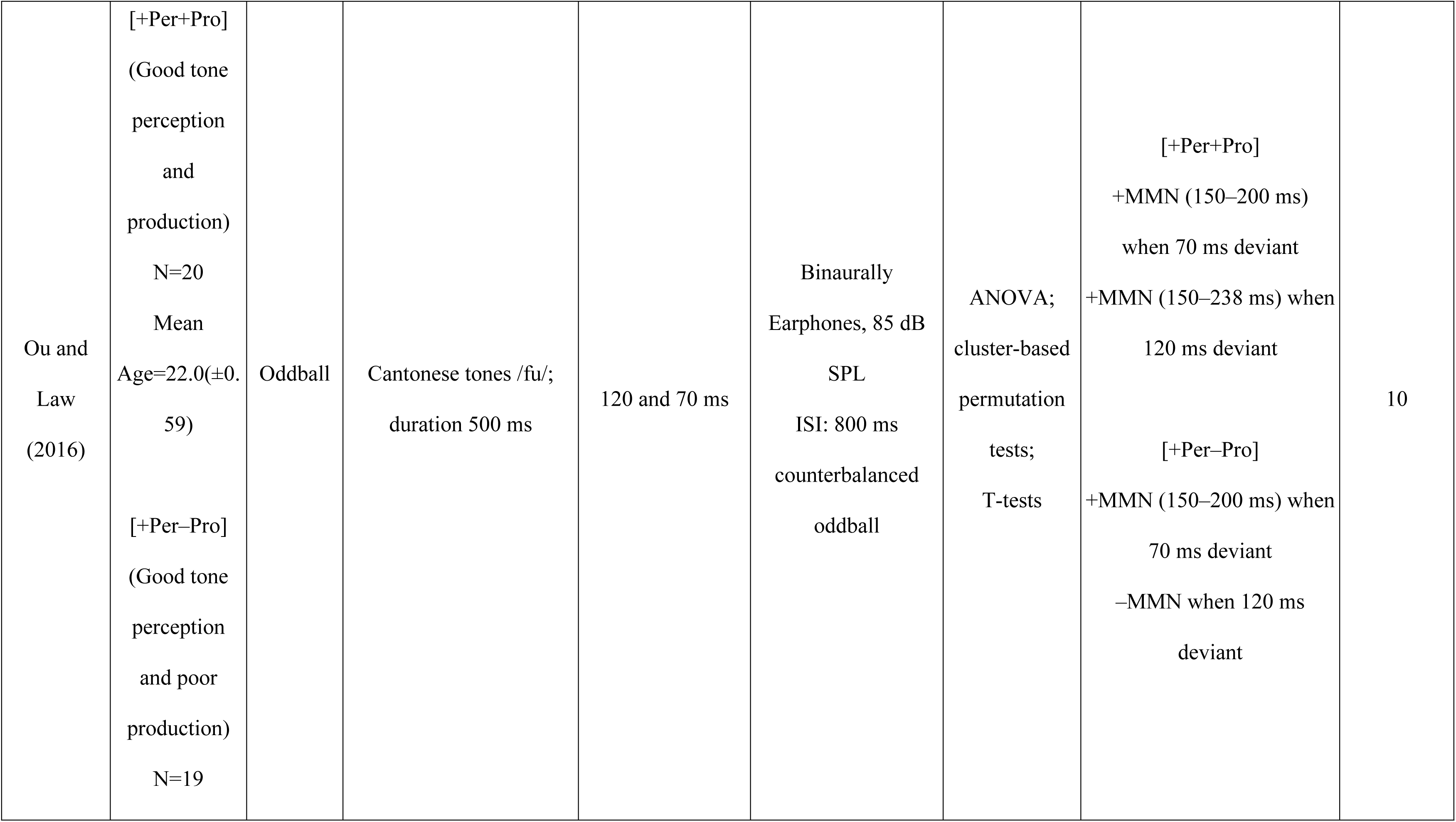
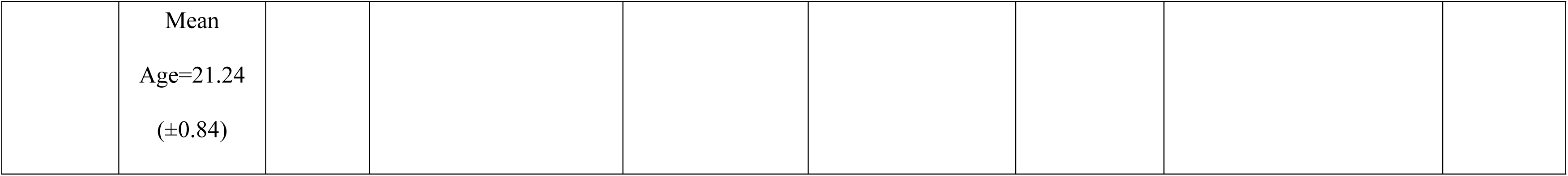
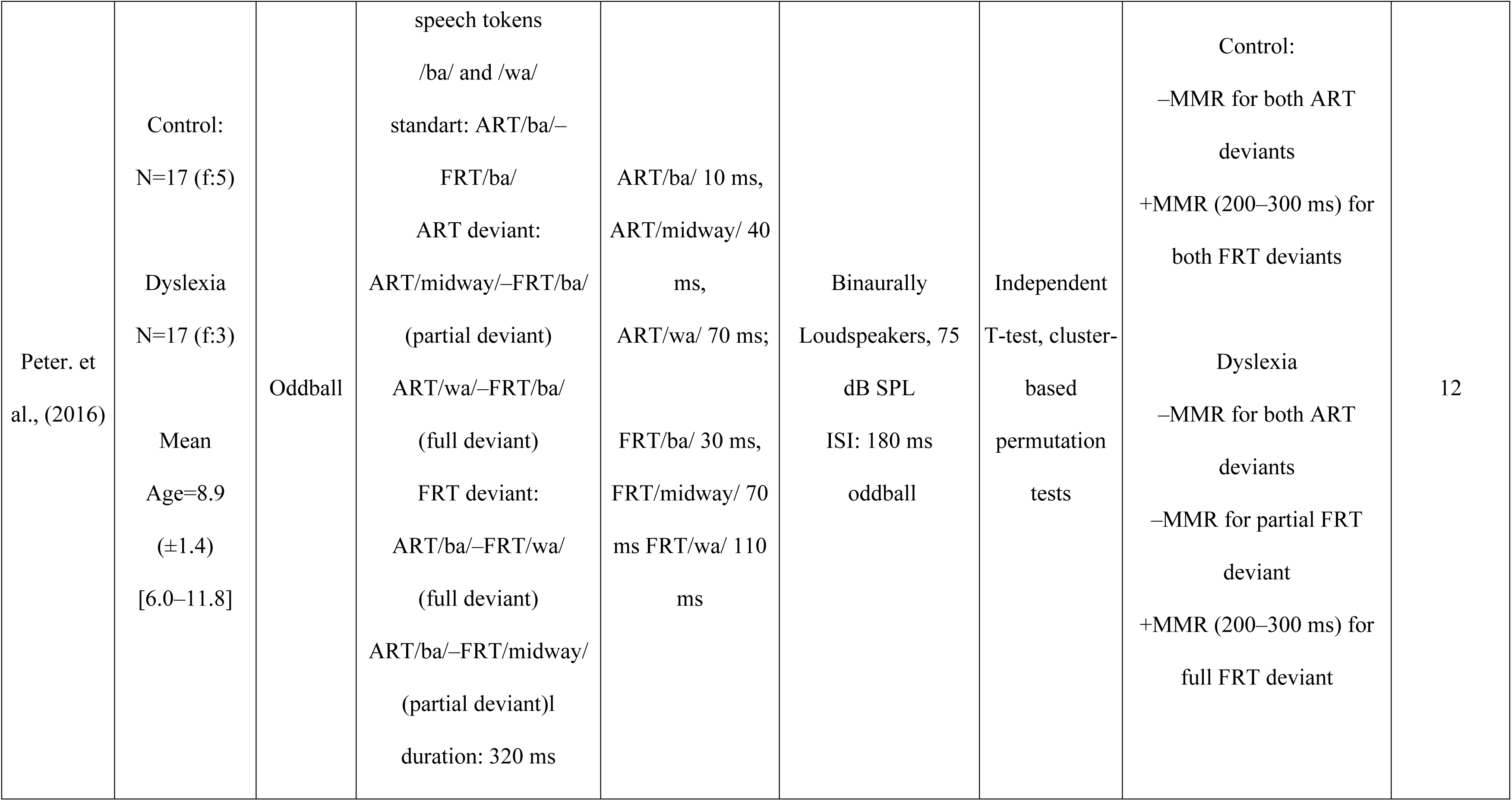
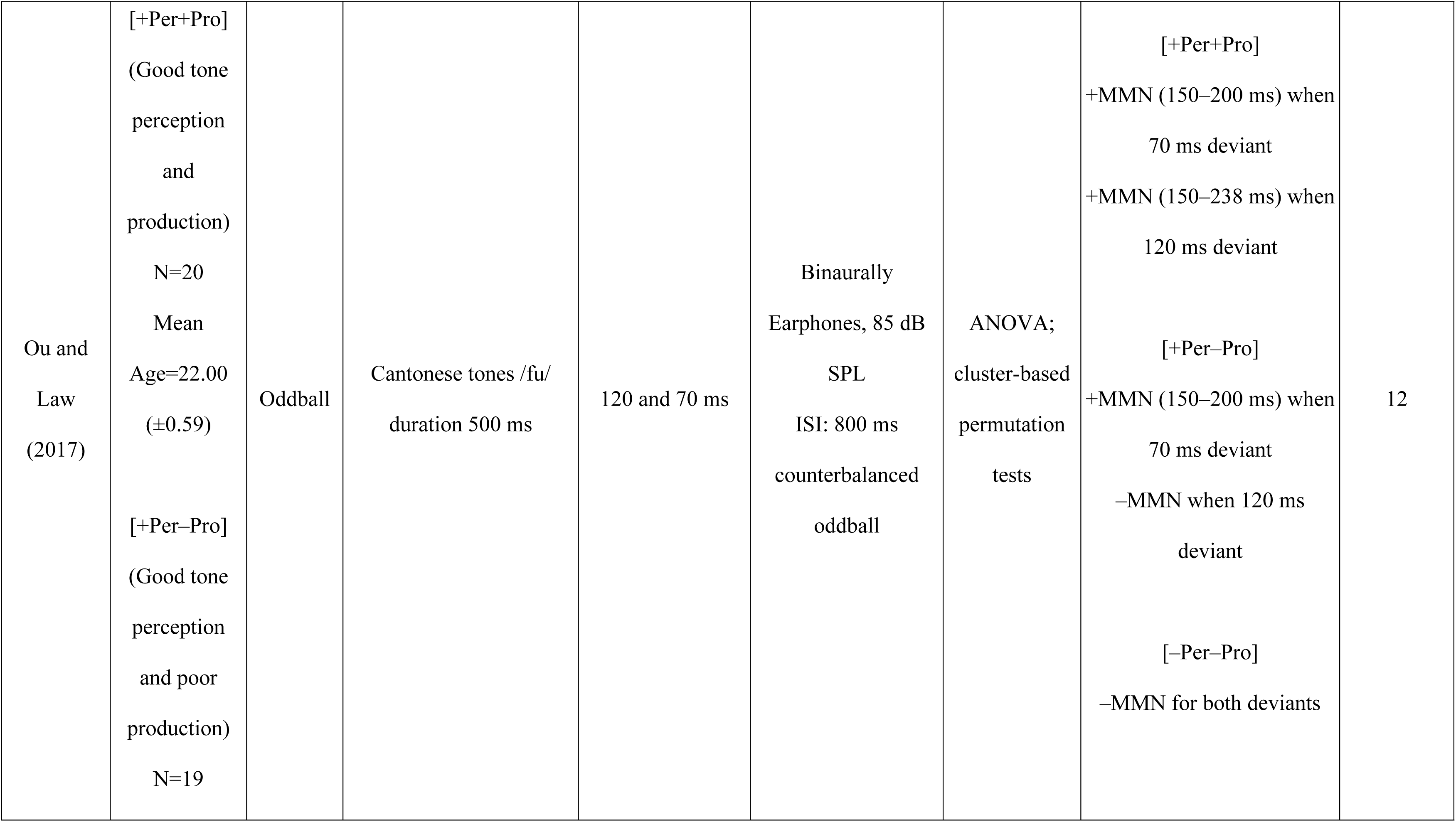
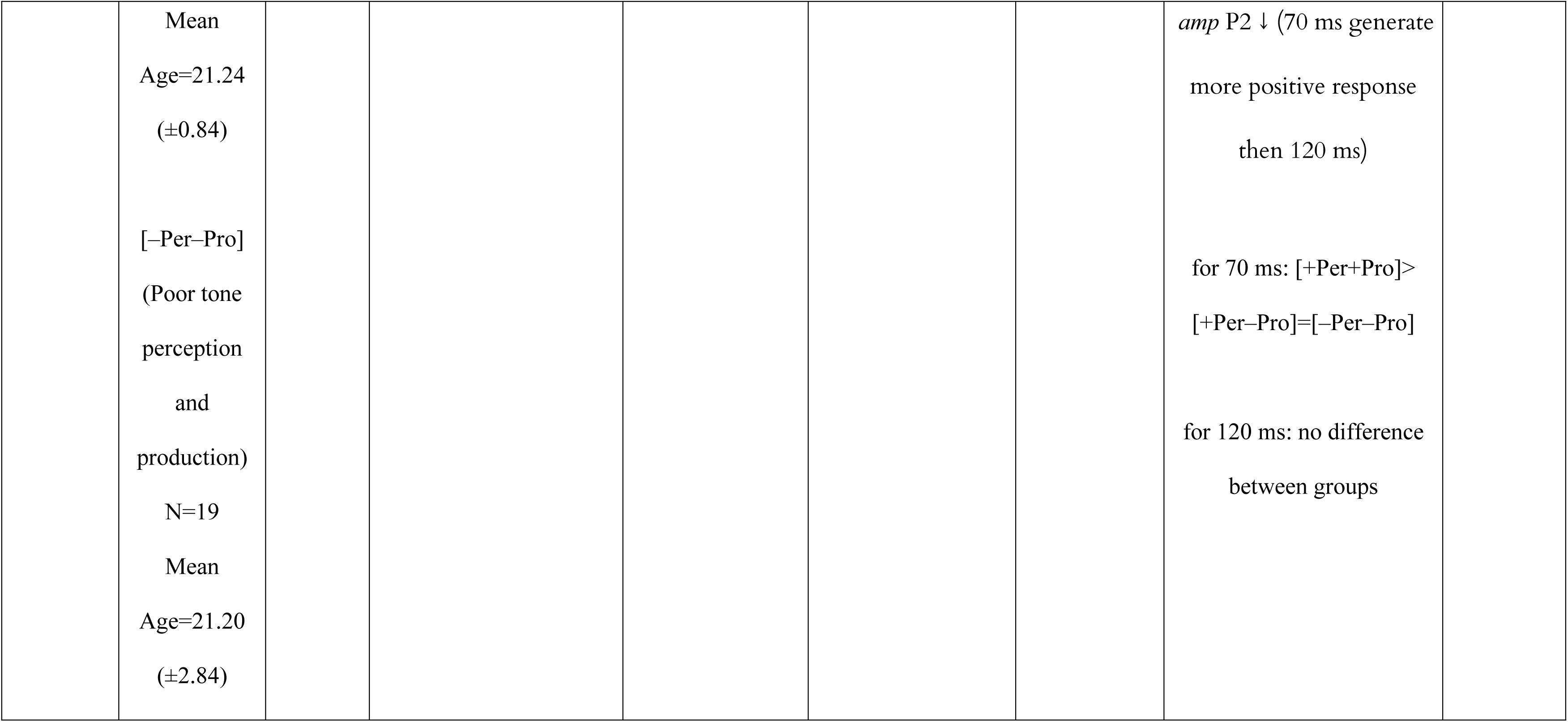
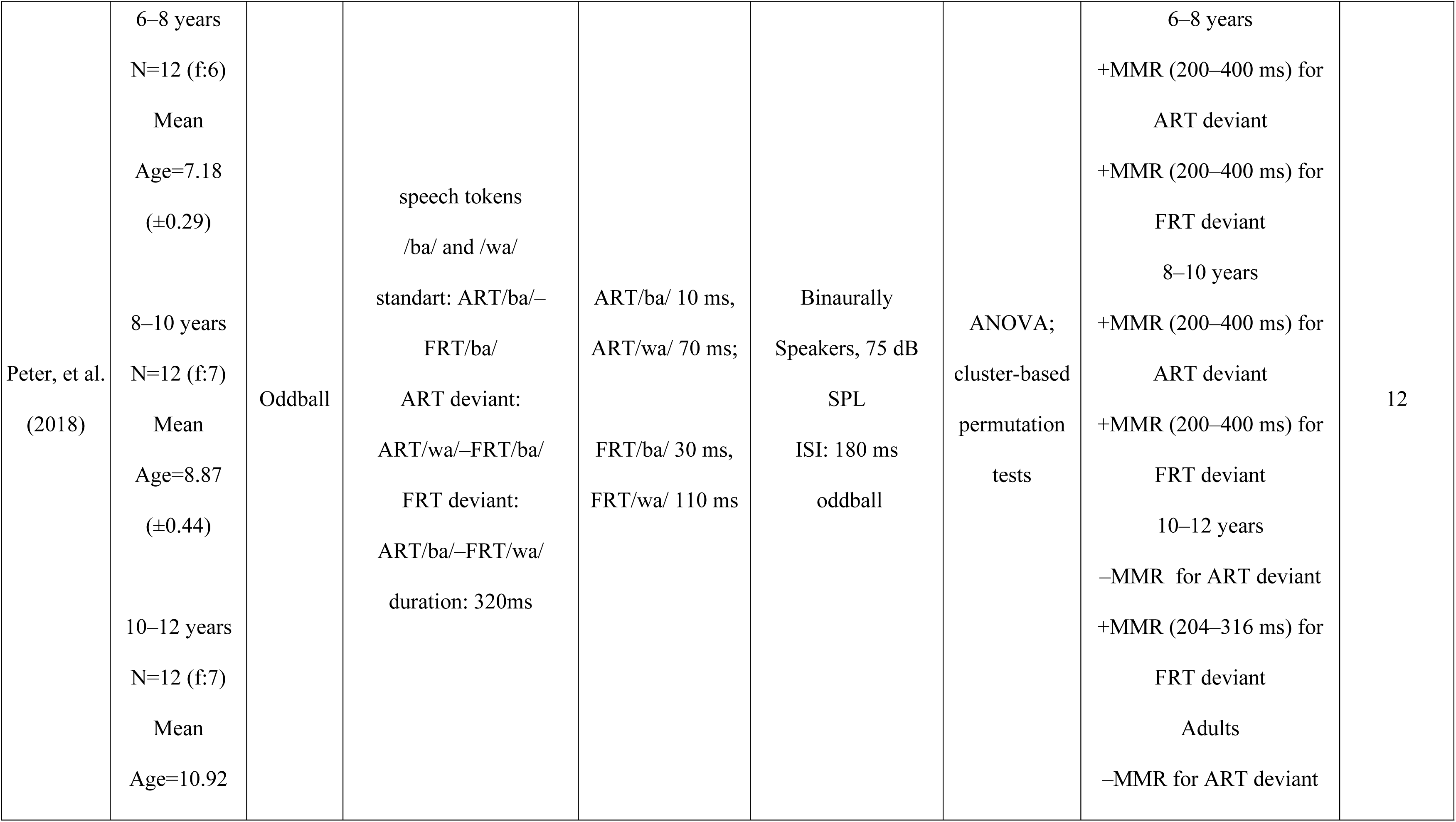
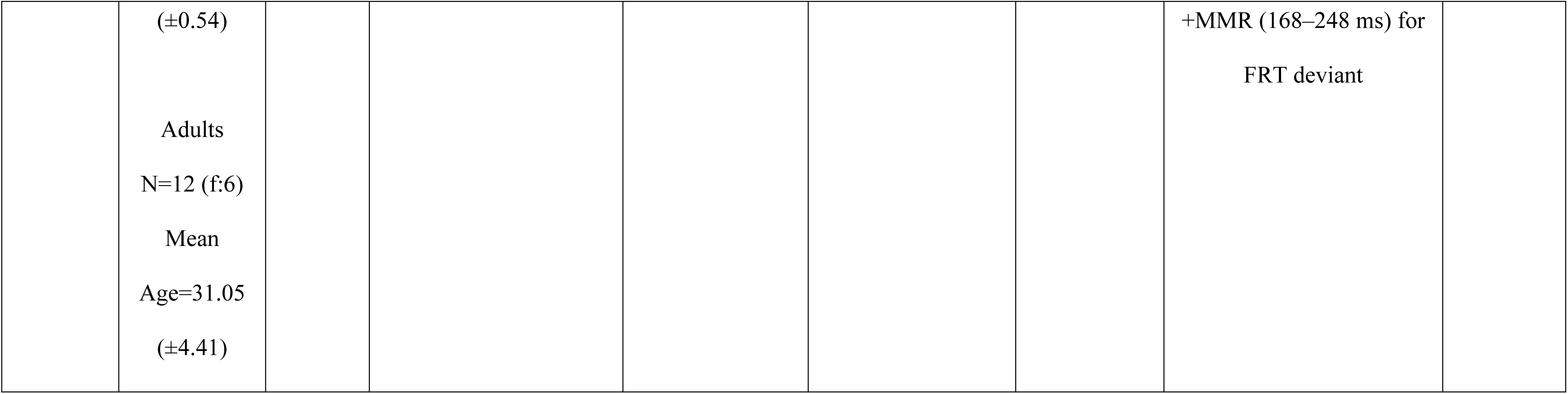
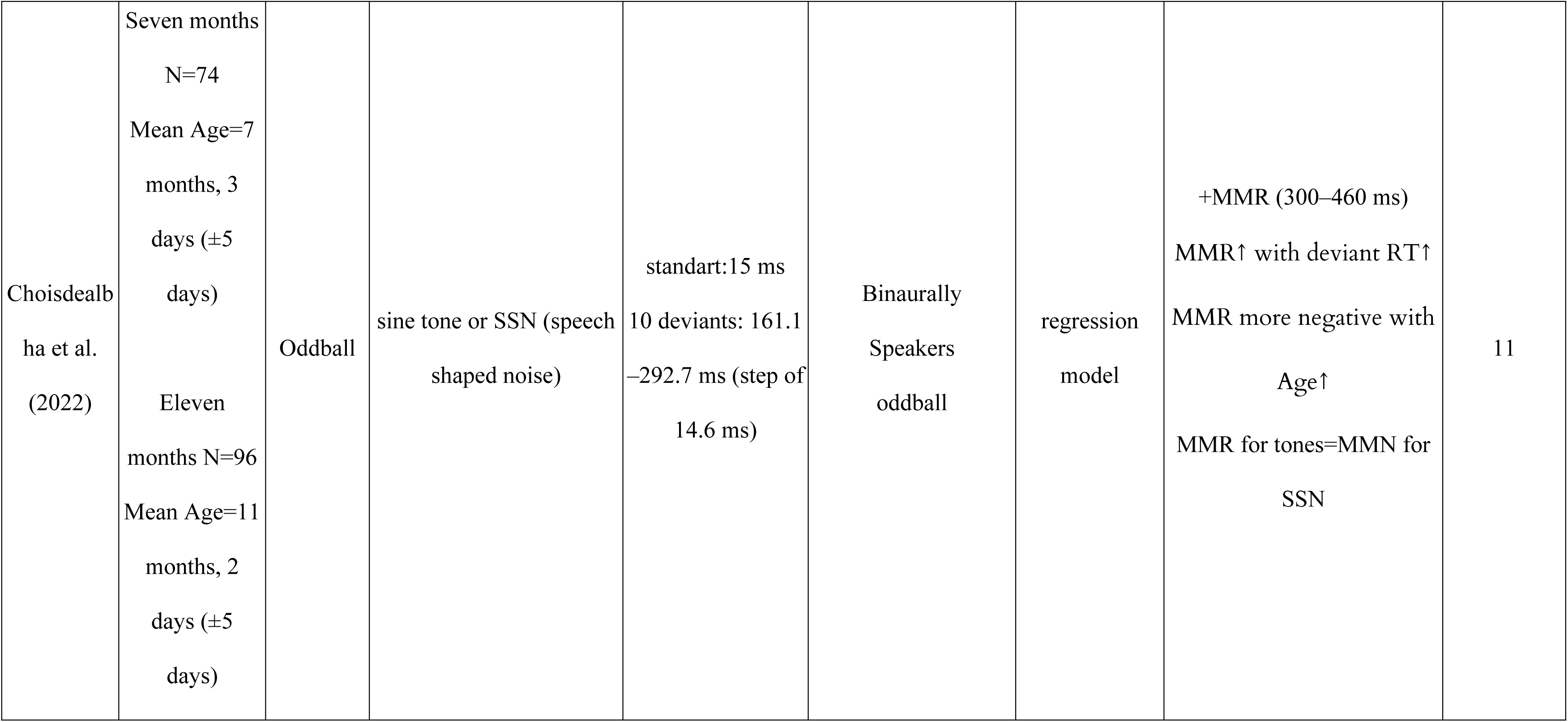
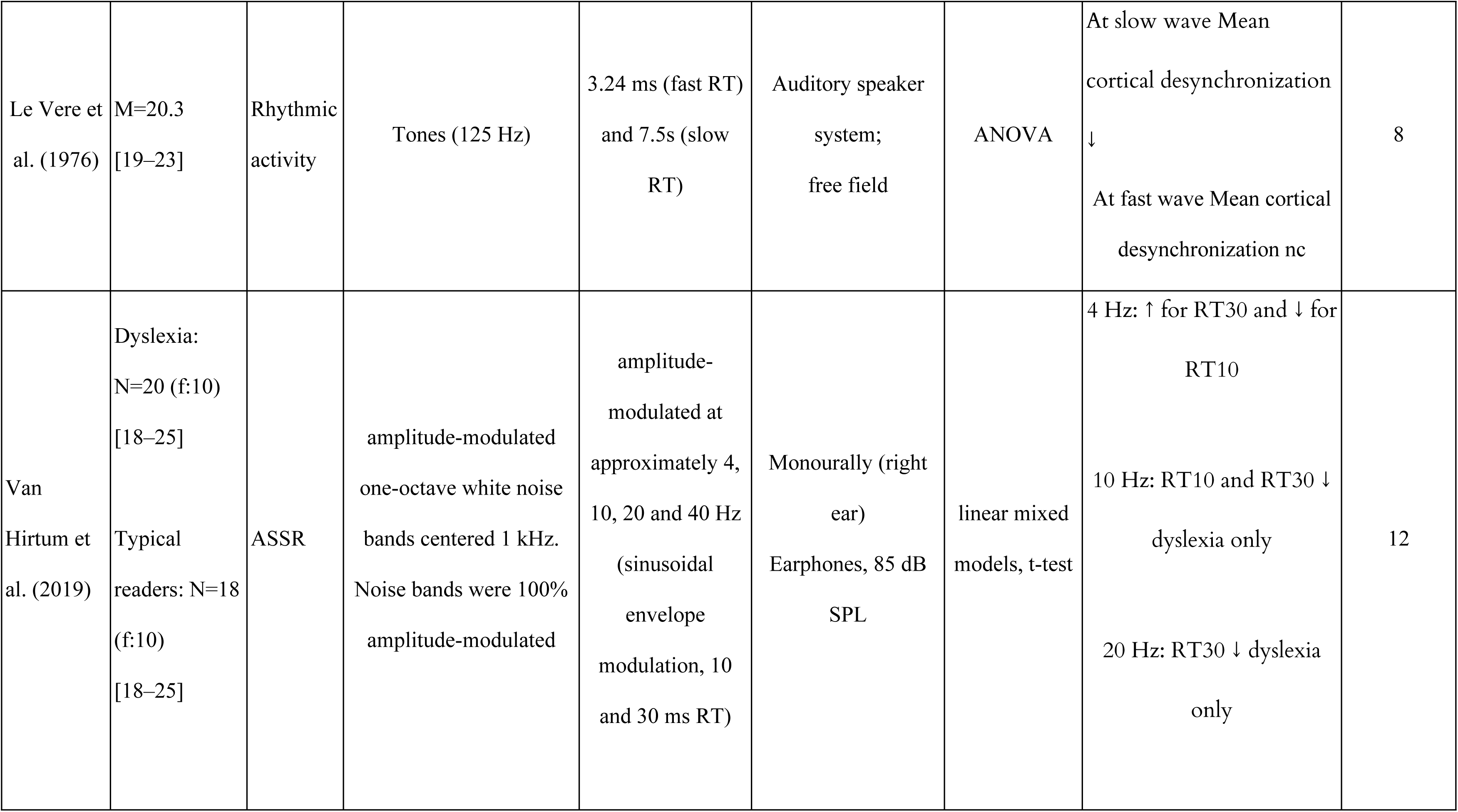
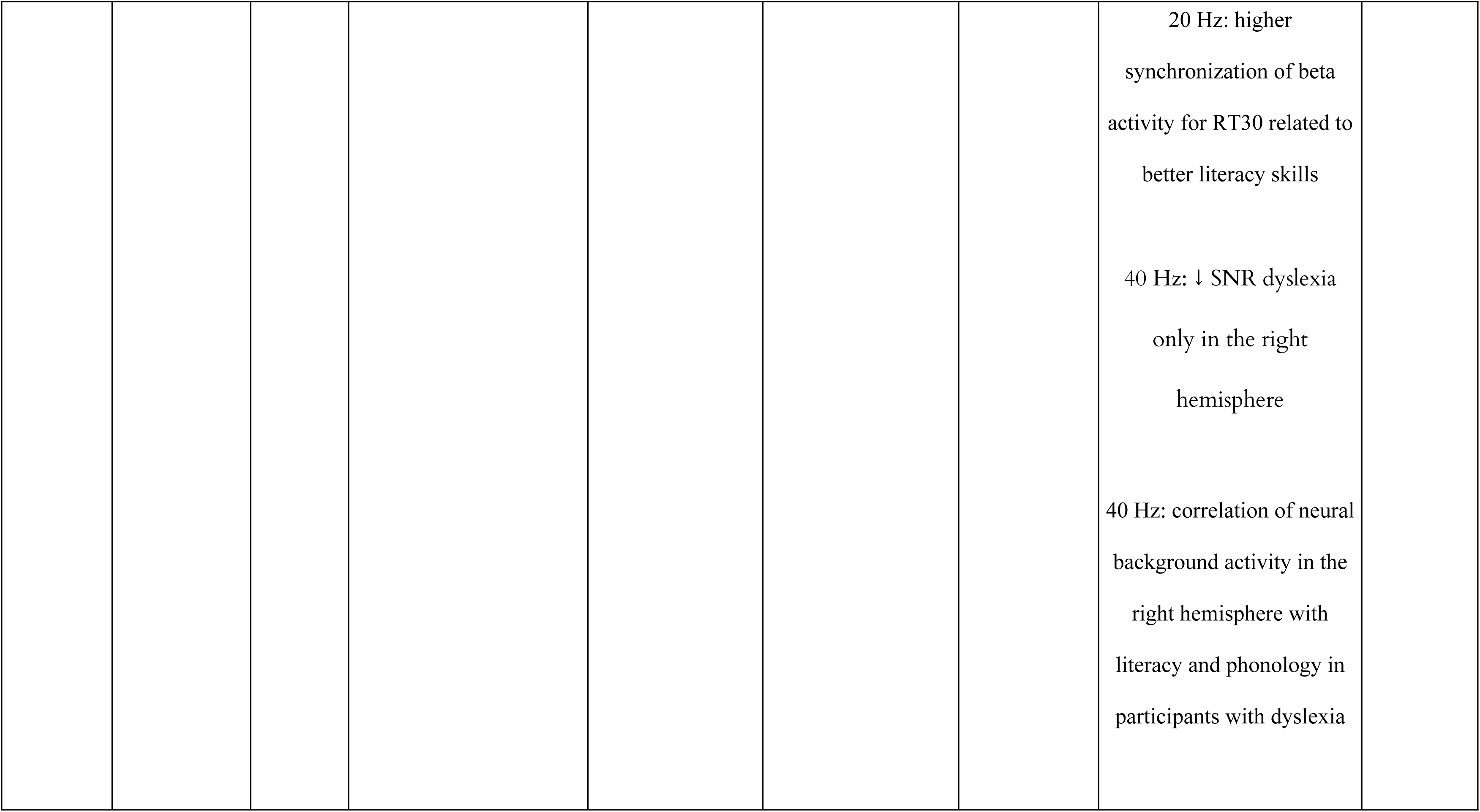
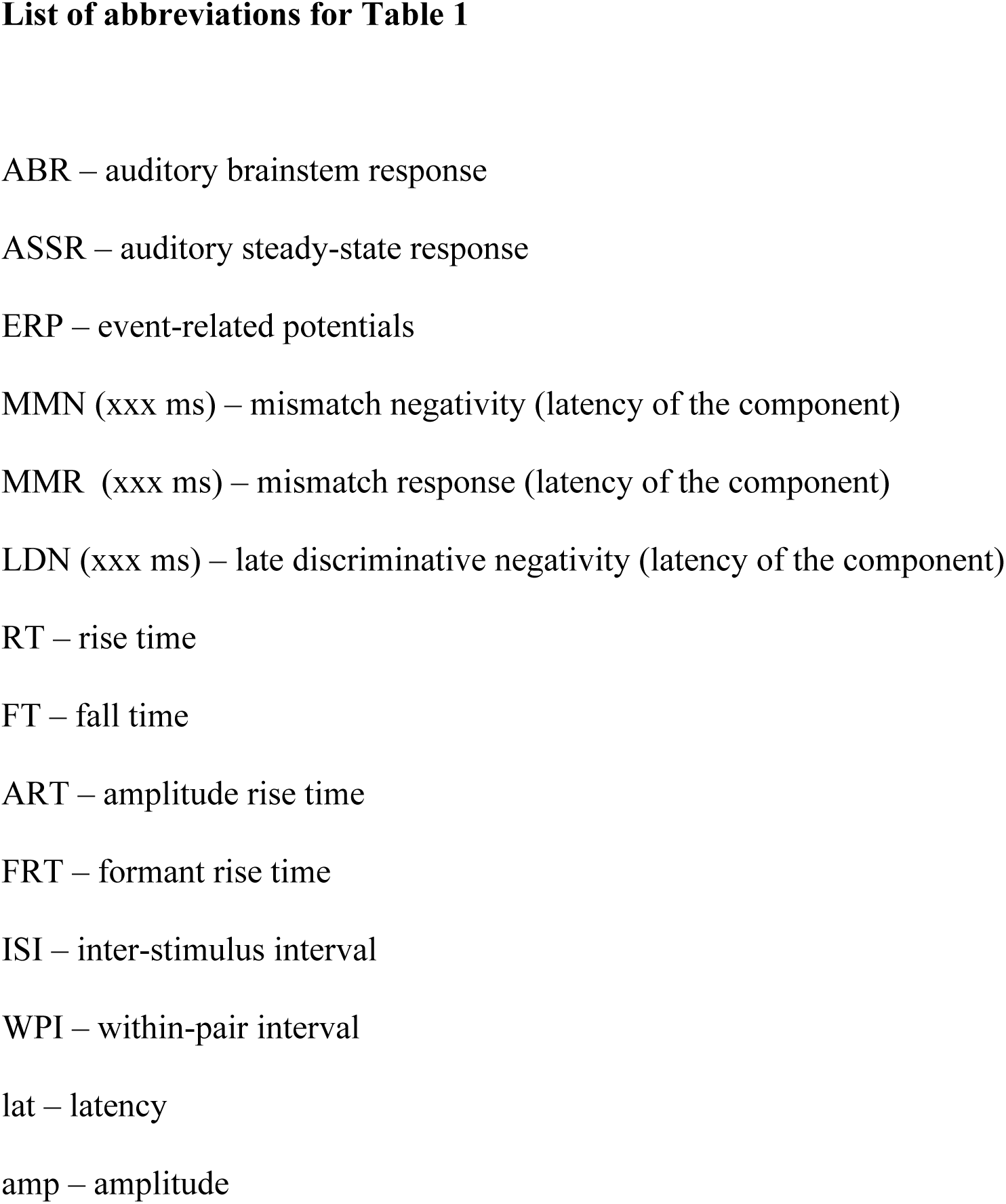
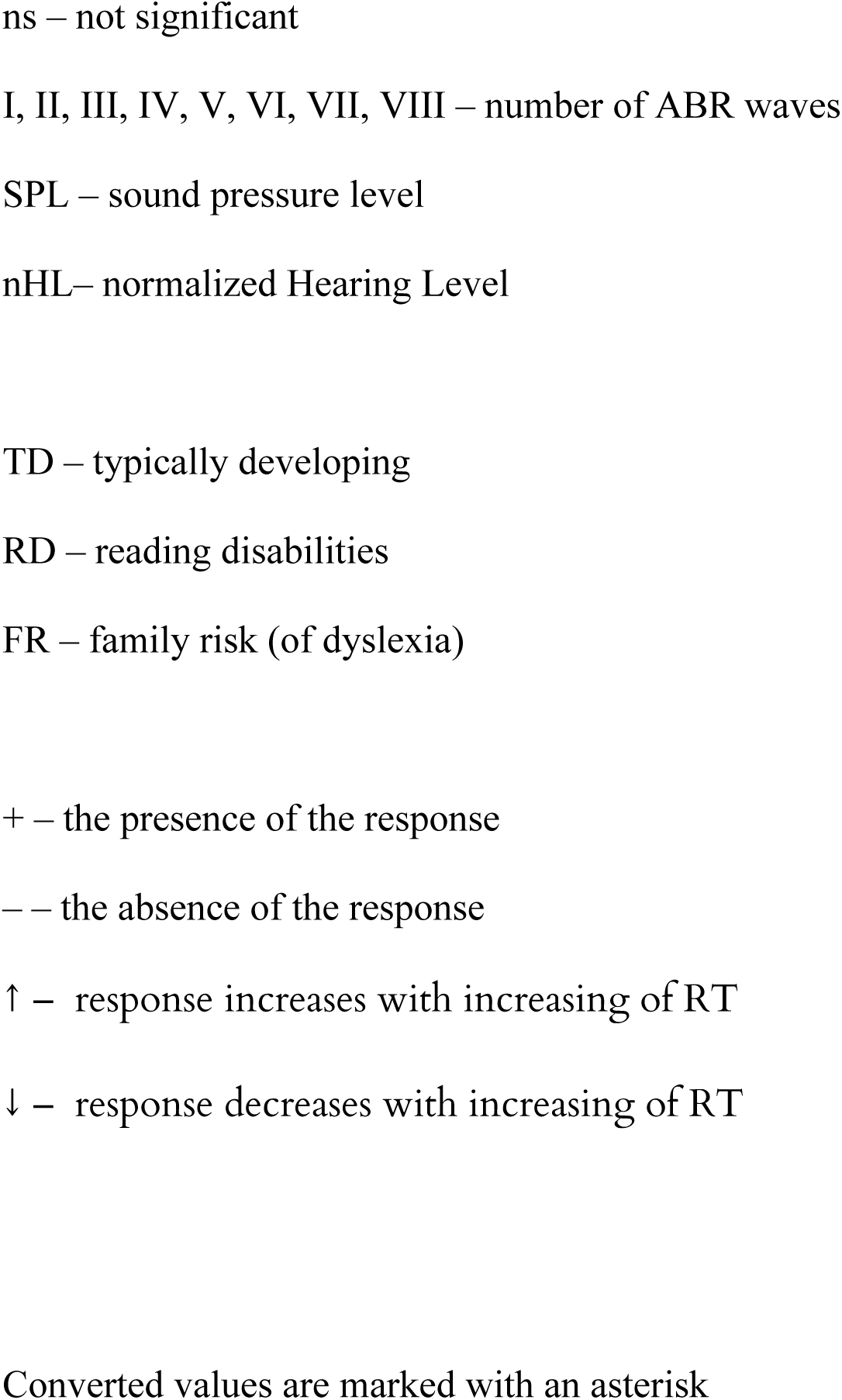
Summary details of included studies. The studies are divided into groups according to the experimental paradigm used, and organized within each group in ascending order of year.

The risk of bias of the included studies was assessed according to the OHAT risk of bias tool [6]. Risk of bias evaluation included 3 sections with several questions inside them: the selection bias, the detection bias, and the “other sources of bias” sections. A table with the results of risk of bias evaluation is provided in the supplementary materials section (<u>Supplementary Material 1</u>).

## Results

A total of 281 articles were identified from all of the used databases after duplicates removal. After abstract and full text screening, 37 articles remained and were included in the review. A flowchart of this selection process is displayed in Fig 1. Summary details for the reviewed studies are presented in Table 1. An updated search conducted on February 15, 2026, identified 13 new records. Titles and abstracts were screened. None of these records met the inclusion criteria for further review (all 13 records were excluded at the title/abstract screening stage, either as irrelevant or as not meeting the inclusion criteria). Therefore, the number of studies included in the review remained unchanged (n=37). The flowchart has been updated accordingly.

Seven articles were excluded from the study due to unavailability of full texts. This was due to the fact that the articles were published a long time ago (1964–1994) and their full texts were not digitized or not available [7–13]. Thirty-seven included studies were divided into groups according to EEG responses studied and organized in the Table 1 and in text according to the latency of the brain responses, from earlier to later evoked potentials. The studies included in the final review ranged in date from 1968 to 2022 (Fig. 2), with an 11–year gap in rise time studies (there were no published studies between 1996 and 2007) and current gap within the last four years.

**Fig. 2.**
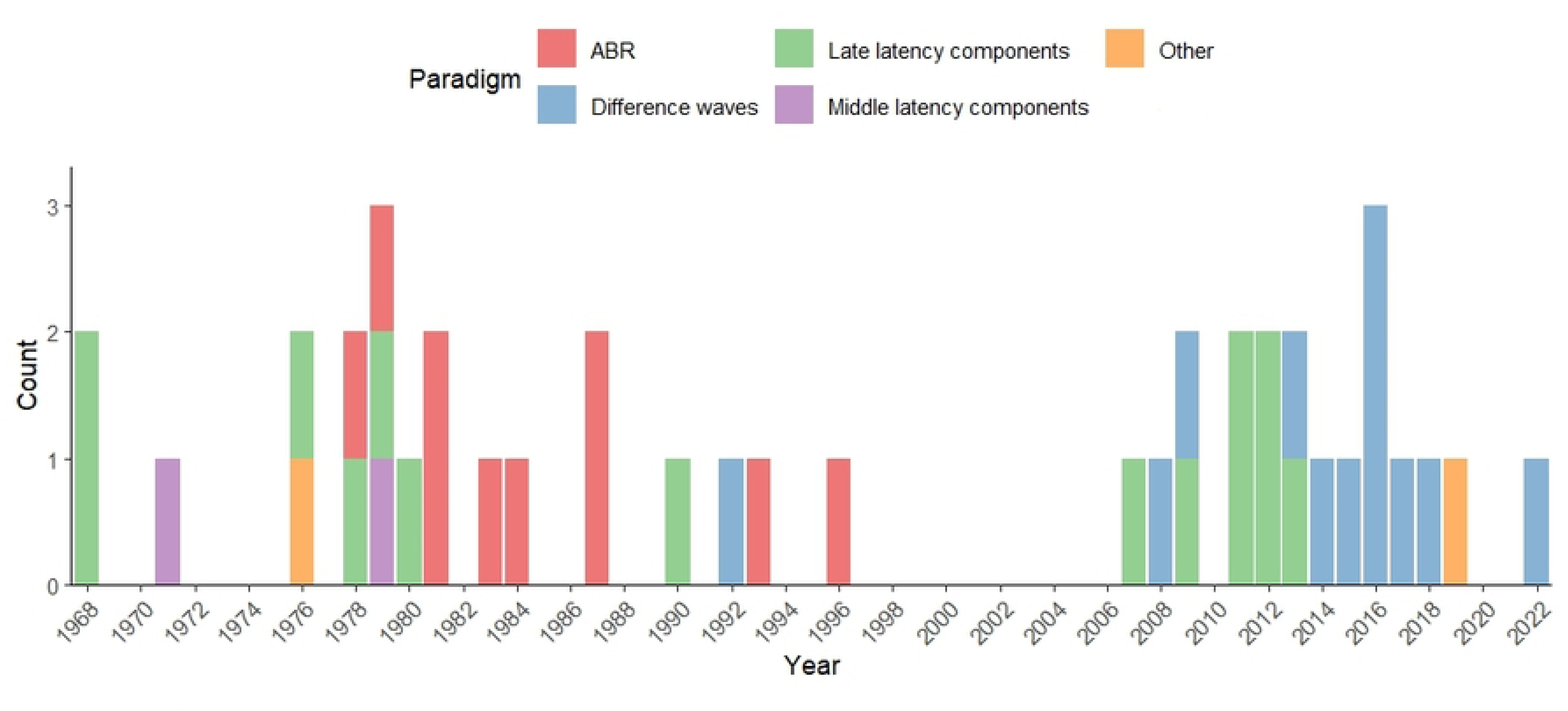
**Histogram of RT studies paradigms over years.**

The main categories into which we classified the studies are: (1) early latency components (auditory brainstem response N=10, (2) middle latency components (Na, Pa, Nb, N=2), (3) late latency components (P1, N1, P2, N2, P3, N=14), (4) difference waves studies (mismatch negativity or mismatch response (MMN and MMR), later positive shift, late discriminative negativity (LDN) – 13 studies and one study on Auditory Steady-State Response (ASSR) and one on mean cortical desynchronization both categorized as Others. Some studies described more than one category of EEG-response and, thus, were reviewed in several sections. Next we describe the results separately for each category.

### Early latency components (auditory brainstem response, ABR)

Auditory brainstem response (ABR) is an early potential that occurs in response to auditory stimuli, reflecting neural activity along the auditory pathways from the inner ear to the brainstem. ABR consists of a series of waves (denoted as I–VII) that appear within the first 10–15 ms after the sound stimulus onset and reflect early auditory processing steps. As ABR occurs within such early latencies, the typical RT studied in this paradigm varies from 0.5 till 5 ms. RT has been studied with ABR for a relatively long time, and no modern studies relevant to the review were found. The most recent study included in this review dates back to 1996. The final review included 10 articles investigating brainstem evoked potentials. The overwhelming majority of studies was conducted on adults, only one of them included 11–13 weeks old infants as one of the experimental groups. However, no difference in the results was found between the adult group and the infant group [14].

The results obtained in studies of RT perception using ABR are quite congruent. One of the main conclusions made by the authors in their articles related to the latency and amplitude of wave V: its latency increases while the amplitude decreases with the RT increase[14–18]. This tendency remains with any type of stimuli (noise or tone bursts, tone pips or clicks).

An increase in latency with an increase in stimulus RT is also observed in other early components of brainstem potentials: from waves I to VII, however, the results depended on stimuli parameters. For example, at 40 dB SPL and for stimuli with the so-called “fast rise time” (10 µs), the latency of waves VI–VII first decreases (between 10 µs and 0.5 ms) and then gradually increases, although this effect was statistically insignificant [19]. In the study of Hecox & Deegan (1983) an increase in the latency of wave V with an increase in RT was also found in most experimental conditions, however, no significant changes in amplitude were found [20]. The study by Suzuki & Horiuchi (1981) was distinguished by its findings of different tendencies depending on stimuli presented. When utilizing 2–kHz tone pips, a decrease in wave V latency with RT increase was observed at certain stimulus intensities (only 30–40 dB nHL). At the same time, a RT change did not significantly affect wave V amplitude. In contrast, with 0.5–kHz tone pips, an increase in stimulus RT correlated with an increase of the mean wave V amplitude and concurrently, the mean wave V latency also exhibited a tendency to increase. However, these results should be treated with caution as they were obtained on a limited sample (N=8, all females) in specific conditions: during sleep induced by administration of pentobarbital calcium before the test [21].

One of the articles reported morphologically similar to onset brainstem response offset brainstem response which occurs by long-duration tone burst (>8 ms) within 8 ms after stimulus offset [22]. Offset ABR was even more affected by stimulus RT than onset ABR. It was demonstrated that both offset and onset wave V amplitude increased and both offset and onset wave V latency decreased (for both 0.5 and 2 kHz stimuli) for 1 ms RT in comparison to 0.5 ms RT. In the block where only two RTs (0.5 vs 5 ms) varied with a constant fall time, the difference were detected only for the offset ABR: offset ABR latency increased for both types of stimuli (0.5 and 2 kHz) and offset ABR amplitude decreased for 2 kHz stimuli with longer RT [22].

Stimulus polarity is a physical characteristic of a stimulus that might influence the RT effect. Stimulus polarity refers to the initial deflection of the transducer diaphragm in relation to the tympanic membrane during stimulus presentation: a rarefaction stimulus causes the earphone diaphragm to move outward initially, resulting in an outward movement of the tympanic membrane, while a condensation polarity stimulus leads to an inward movement of the diaphragm and consequently an inward movement of the tympanic membrane [23]. Most ABR studies included in the current review used alternating stimulus polarity (rarefaction and condensation), thereby cancelling out the polarity-specific effect. Additionally, some investigations within the ABR-field have explicitly considered stimulus polarity as one of the influential acoustic parameters, alongside other stimulus characteristics. Amplitude values of main ABR components in study of Salt & Thornton depended on polarity of stimuli presented: direction of amplitude changes was different in rarefaction vs condensation stimuli. In particular, I, II, III, VI amplitudes in response to rarefaction stimuli tended to be lower with RT increase and vice versa on condensation stimuli. Amplitude of peak V, on the contrary, arose with RT in response to rarefaction stimuli and lowered with RT in response to condensation [24]. However, no statistical testing was applied. Besides, the other study which used clicks of different polarity as stimuli [16] showed the same direction of amplitude changes in both stimuli types. However, the latencies in all RT conditions were longer for condensation slope stimuli, and amplitudes were contrariwise higher for rarefaction stimuli [16]. In the case of short stimuli with a RT of 170-580 µs, there is a linear tendency towards an increase in the latency of the main components (waves I–VI) with an increase in RT for stimuli of both polarities [24].

### Middle latency components

Middle latency components are considered to be driven by the primary auditory cortex with a substantial contribution of thalamic-cortical pathways and elicited from 10 to 80 ms after stimulus onset [25]. In this group of components, several successive peaks are distinguished (P0, Na, Pa, Nb, Pb). Two studies[26,27] demonstrate the modulation of middle latency components of evoked potentials by RT duration in a group of neurotypical adults. Tone bursts were used as stimuli. In both of these studies short RT durations were used (up to 25 ms). In [27], peak-to-peak amplitude of Na–Pa, Pa–Nb components decreased with RT increased across all conditions, while peak-to-peak amplitude of P0–Na and Nb–Pb components decreased only in the longest RT condition (25 ms). Notably, no statistical analyses were used in this study, so the presented results should be considered with caution. In a later study by Kodera et al. (1979) [26], the decrease in Na–Pa, Pa–Nb peak-to-peak amplitudes with RT increased was verified statistically. This article also showed a delay in the Na and Pa components latencies with RTs prolongation.

### P1

The P1 component is a positive peak in the auditory ERP, occurring approximately 50–100 ms post-stimulus in adults (>100 ms in children), and originating primarily in the primary and secondary auditory cortex. In neurophysiological research, P1 is optimally recorded from central midline electrodes, particularly Cz referenced to mastoids or earlobes, and serves as a biomarker for auditory cortical development and central auditory pathway integrity [28]. P1 has rarely been an object of interest in RT studies as it is not very prominent in adults: one study in children and one in adults. Stefanics and colleagues [29] examined P1 modulation by RT duration in dyslexia and typically reading children (aged 8–10 years) in longitudinal study that includes oddball and block conditions. Decrease in P1 amplitude with RT increase was observed only in the group of participants with dyslexia, but not in the control group, where P1 amplitudes were equivalent across RT conditions. This effect was specific only for oddball conditions; no significant effects were observed for P1 amplitude in the blocked condition. Authors attribute this difference to greater responsiveness of the P1 component in oddball condition, which was related to different probability of occurrence of the stimuli. Both groups demonstrate prolongation in P1 latency with RT increasing in all conditions. The same prolongation was observed in Kodera et al. (1979b) [26]. In this study P1 was considered as part of P1–N1 peak-to-peak amplitude, which demonstrates significant decrease with RT prolongation. In Skinner and Jones (1968) [30] work no significant effects for P1 latency were observed.

### N1

The N1 (or N100) is a prominent negative-going component of the auditory ERP waveform that follows P1 and typically peaks approximately 80–120 milliseconds after stimulus onset. It is primarily generated in the auditory cortex, including areas within the superior temporal gyrus (such as Heschl’s gyrus and the planum temporale). In EEG recordings, the N1 component typically exhibits maximal amplitude over fronto-central scalp locations and is therefore commonly observed and analyzed at midline fronto-central electrodes such as Fz, FCz, and Cz [31].

Of all the articles included in the review, thirteen examined the effects of RT on the N1 component. According to reviewed studies, reduction in N1 amplitude with RT prolongation is a common pattern for neurotypical adults. In [32] N1 amplitude linearly decreases across RT prolongation from 3 to 45 ms, but only in the first of two runs of the paradigm. Thomson and colleagues (2009) [33] shows that N1 amplitude also decreases in longer RT condition, such as 50 and 185 ms. These results are confirmed in the additional part of that study, where stimuli with different RT were equalized in intensity. Similar reduction of N1 amplitude in longer RT conditions were obtained for groups of typically reading adults [34] and children [35]. In one of the studies[26], N1 was assessed as P1–N1 peak-to-peak amplitude, which also demonstrates significant decrease with RT prolongation from 5 to 20 ms.

Three studies reported significant enhancement of N1 latency with RT increase[26,34,36]. A similar result was shown in earlier papers [37,38], but was not confirmed by statistical analysis. In the study by Eswar and colleagues[39], the difference in N1 latency between RT conditions was not significant, which probably related to minor differences between RT conditions used in this work (7.5 vs 20 ms). However, even larger contrast (such as 2.5 vs 50 ms) was also insignificant for 1000 Hz tone burst and white noise burst [40].

Two studies examined N1 modulation by RT in a group with speech impairment. In Hämäläinen et al., (2007) [35] in a pair of stimuli with a short within-pair interval, the amplitude decreases at a longer RT condition (80 ms), but only in typically developing (TD) children. In the group of children with reading disabilities, the N1 amplitude was equal across RT conditions. It is important to note that stimuli differed not only in RT but also in frequency and statistical comparison of different RT conditions was not performed. The more recent work of this group [34] with stimuli of the equal frequencies showed decrease in N1 amplitude and prolongation of its latency with RT increase in both typical reading and dyslexic adults groups.

### P2

The P2 component is a positive deflection in ERP that follows the N1 component and occurs approximately 150–200 ms post-stimulus. It is generated in the auditory association cortex and surrounding areas and thus, is typically recorded from central and frontocentral electrode sites (Cz, FCz) and suggests to reflect higher-order sound processing, auditory feature integration, and early attentional mechanisms [31].

Nine of the papers included in the review consider RT effects associated with the P2 component. Most works estimated P2 effects as related N1–P2 peak-to-peak amplitudes. N1–P2 peak-to-peak amplitude was considered in seven studies[17,36,37,39–42]. N1–P2 amplitude tends to decrease with prolongation of the RT. In Onishi & Davis work [37] RT changes did not affect the N1–P2 amplitude when RTs were less than 30 ms. Decrease in N1–P2 amplitude was observed only as RT is increased beyond about 30 ms, but no statistical analysis was performed. In Kodera et al. (1979) [26] no significant difference was shown for N1–P2, but RT durations used in this study were quite short (5–20 ms), which probably not enough for N1–P2 modulation. A similar result appears in a more recent study[39], where no difference was found between 7.5 and 20 ms RT conditions for 1000 Hz tone burst. [40] observed significant decreases in P2 for 50 ms RT condition compared to 2.5 ms RT condition for 1000 Hz tone burst, but not for white-noise burst. In the study by Elfner et al. [42], decrease in N1–P2 amplitude was shown for prolongation RT from 50 ms to 150 ms and 500 ms, although continued prolongation from 500 to 1500 and 4960 ms of the RT duration did not produce a change in N1–P2 amplitude. However, further statistical analysis in the Experiment 2 of this study showed that the observed RTs prolongation effects are only at the level of tendency and are not statistically significant. Two studies[36,41] used more natural types of stimuli, such as speech tokens. N1–P2 amplitude decreased in longer RT (such as 79.1 ms in Easwar et al.; in Carpenter & Shanin exact RT not reported) in both studies. However, the effect observed in adults and 6-years old children, was not present in early childhood (4–5 years) [41].

Two studies [32,34] considered P2 as an absolute baseline-to-peak value, not as a peak-to-peak amplitude. Putnam & Roth [32] showed linear decrease of P2 amplitude (considered as P190 component) with RT prolongation that included RT ranged from 3 to 45 ms. This result was confirmed in the study by Hämäläinen and colleagues (2011) [34] both for control and dyslexic adults groups with RT used from 10 to 120 ms.

As in case of N1, P2 latency demonstrated delay with RT prolongation[26,36]. At first, this effect was described in [38] as shifting of slow V potential (N1–P2 peak-to-peak component), but was not statistically validated. Later it was confirmed statistically[17,36]. Nonetheless, in some papers, the differences in P2 latency between different RT conditions were not significant[34,39].

### T–complex

Two of the studies included in the review examined RT effects not only in the central but also in the temporal channels. Näätänen & Picton [43] define a projection of the P1-N1-P2 components to the temporal region as T-complex. The T-complex includes three peaks: N1a (or Na) (75–95 ms), N1b (or Ta) (100–115 ms), and N1c (or Tb) (130–170 ms). In the study of Hämäläinen et al. (2011) that included neurotypical adults and adults with dyslexia, Na and Tа amplitude did not change between RT conditions in both groups, while amplitude of Tb decreased with RT prolongation in a group of adults with dyslexia, but not in the control group of neurotypical adults. There was also prolongation in Na component latency in both groups. In a group of children (7–10 years) Stefanics and colleagues [29] also showed decrease of N1c (at FT7–FT8 sites) amplitude with long (90 ms) RT in oddball condition, but in the block condition RT effects were observed only in dyslexic group. No RT effects on N1c latency were observed.

### N2

The N2 component is a negative deflection in the auditory ERP that follows the P2 component and occurs approximately 200–250 ms post-stimulus onset. It originates primarily from frontal cortical regions including the anterior cingulate cortex, which is typically recorded at frontocentral electrode sites (Fz, FCz) [44]. Only one study [29] considered N2 as a target for the RT effects in children from 7 to 11 years. Similar to the other components, N2 demonstrated decrease in amplitude and prolongation of latency with the RT increase. One more study by Skinner & Jones [30] considered N2 as part of the P1-N2 complex, where the difference between P1 and N2 components was taken as the amplitude of the auditory evoked response. It was shown that P1-N2 decreases with RT increase. N2 latency did not show RT effects, but statistical analyses were not performed. Even though the effects of RT on the N2 component were not considered in other works, we can assume decrease in amplitude and latency delay when the RT is prolonged by looking at the figures[34,36].

### P3

The P3 (or P300) component is a prominent positive wave occurring approximately 250–500 ms following auditory stimulus presentation, generated by distributed neural networks involving frontal, temporal and parietal cortical regions. P3 is reliably recorded at midline electrodes (Fz, Cz, Pz) with maximum amplitude typically at Pz [45]. One study [32] considered the RT effects for late positive component (P300). Amplitude of P300 (at Pz electrodes) significantly decreased with RT prolongation from 3 to 45 ms, however the RT prolongation was also accompanied with changes of the total duration of stimuli that can also influence the observed change [32].

### Difference waves (MMN, MMR, Late positive shift, LDN)

Many studies investigating RT processing have employed the oddball paradigm, where deviant stimuli with different RTs are embedded within streams of standard stimuli with fixed RTs [33,46–48]. These deviant stimuli typically elicit a mismatch negativity (MMN) or mismatch response (MMR). MMN is obtained as a difference between ERP in response to standard (frequent) stimuli and ERP in response to deviant (rare) stimuli. Sometimes, especially in children, the response to a different stimulus may appear as a more late positive component on a difference wave, which is known as a Mismatch response (MMR). Mismatch Response (MMR) refers to the corresponding response in infants and young children, which can be positive (pMMR) or negative (nMMR) in polarity, with a broader scalp distribution and more variable latency than adult MMN, reflecting the developing auditory system [49]. Twelve of the reviewed papers examined sensitivity to RT changes using this paradigm, with the MMN component serving as the primary measure of distinguishability between standard and deviant stimuli. Two studies considered also late discriminative negativity (LDN), a later (from 400 ms post-stimulus) component at the difference wave evoked by a deviant stimulus and also suggested to be localized in associative areas of the auditory cortex. Although originally LDN was called the late MMN [50,51], included studies considered it as a separate component to highlight the difference in time of occurrence and to separate the RT effects.

The first study reported MMN (in time window 150–250 ms) elicited by RT deviants in neurotypical adults, used short RT durations (24 ms in standard and 2 ms in deviant) [52], also late positive shift in differenсе waves (at time window at 280–500 ms) was described in this work. Later study of neurotypical adults [33] used two RT durations (15 and 185 ms). Each type of stimulus was used as standards and as deviants in different blocks. Both types of deviant elicited MMN response (in time window 150–250 ms) in frontal and temporal areas. In the second part of this study, four types of stimuli were used: one with long RT (50 ms), two with RT 15 ms (one of them was intensity matched to stimulus with 50 ms RT, and other was not), and one with short RT stimuli that was equivalent in intensity to long RT stimulus (intensity matched deviant). This contrast was introduced to separate the influences of intensity and RT effects, as changing RT inevitably influences intensity especially with long RT. When 50 ms RT stimuli were used as standard intensity-not-matched deviant with shorter RT elicited a MMN response at time window 175–275 ms (but not significant), while intensity-matched deviant did not elicit any MMN at all. However, in the exchanged condition where intensity-matched 15 ms RT stimuli were used as standards and 50 ms RT stimuli were used as deviants, a mismatch response was observed in the time window of 175–275 ms even for intensity-matched condition. According to these results, authors suggested that MMN is sensitive to the “content” of the change (RT) but not purely to the amount of stimulus energy (intensity). Meanwhile the context effect, i.e. the observation of prominent MMN only for conditions with longer RT for the deviant than the standard, was left unexplained.

Controversial results were obtained concerning the MMN to RT deviant as neuromarker for reading problems. The first study [53] considered MMN in response to deviants with short RTs (10 vs 130 ms RT in standard stimuli) in typical reading children and children with reading disabilities (RD) (8–10 years). The study included two conditions with different intervals between stimuli in the presented pair. The authors chose a varying Within-Pair Interval (WPI) design to investigate if children with reading problems exhibit deficits in rapid auditory processing, expecting larger differences with short WPIs, and also to determine if they process rise times differently at an early stage, regardless of the WPI length. This approach allowed them to examine how both rapid presentation rates and rise time variations influence early neural responses in children with and without reading difficulties. Each trial presented a pair of tones under two timing conditions: a short gap condition with 10 ms between tones (within-pair interval, WPI) and a longer gap condition with 255 ms between tones. The experimental design included 80% standard pairs (where both tones had expected properties) and 20% deviant pairs. In deviant pairs, the second tone differed either in RT (10% of trials, 10 ms instead of 130 ms) or in pitch (10% of trials, 750 Hz instead of 500 Hz). The first tone remained consistent across all pairs (100 ms duration, 80 ms rise time, 300 Hz frequency), while the second tone in standard pairs had 150 ms duration, 130 ms RT, and 500 Hz frequency. All tone pairs were separated by a consistent 610 ms interval (ISI), regardless of condition.While in the longer interval condition (255 ms), MMN response (at latency 119 ms from the deviancy onset) was larger than in the shorter interval (10 ms) in both groups, children with RD demonstrated a larger MMN than typically developing (TD) children only in this longer interval condition. This study also considered LDN. In conditions with a short (10 ms) interval after the previous stimulus, LDN was not presented. In long within-pair intervals (255 ms) conditions, an LDN response was observed at 375–645 ms. In this condition, the control group showed larger LDN than children with RD.

A traditional non-paired oddball design was employed by Plakas et al. (2013) [47] in their study of younger children (3 years old). A negative mismatch response at 346 ms for longer RT deviants (90 vs 15 ms RT in standard stimuli) was observed in the control group but not in people with dyslexia and typical reading, but with familial risk groups. The study by Hakvoort and colleagues (Hakvoort et al., 2015) involves similar groups of children, but the age of the participants was older (11 years). MMN and LDN, responses peaking around 260 ms and 420 ms respectively to RT deviants with longer RT (90, 180 or 270 vs 15 ms RT in standard stimuli) were observed for all groups (typically developing children, children with dyslexia, and children with familial risk of dyslexia) and did not differ between groups. The influence of different RT durations of deviant stimuli was not considered in this study. The interstimulus interval in this study was close to the long within-pair intervals condition in the Hämäläinen et al. (2008) study (250 ms). These findings suggest that the presence and magnitude of the LDN component may depend on the timing between stimuli and vary across populations with reading difficulties.

A number of studies have used speech stimuli [4,54–56]. In these studies, the amplitude RT (ART) changes are usually contrasted with the formant RT (FRT) changes. Syllables /ba/ and /wa/ are commonly used stimuli in this type of study. Syllable /ba/ has a short ART (10 ms) and FRT (30 ms), /wa/ has more prolonged ART (70 ms) and FRT (110 ms). The third synthetic token usually used in this approach combines a /ba/-like FRT (30 ms) with a /wa/-like ART (70 ms). The Moberly et al. study [4] found that while neurotypical adults show MMN response between 150–350 ms for changes in both ART and FRT deviants, the MMN is significantly larger and more consistently observed across individuals for FRT deviants. In a further study of this research group [54], similar experimental paradigms were used for adults with cochlear implant sample. MMN response was observed for ART and FRT deviants and did not differ between contrasts. Peter et al. [55] implicated FRT and ART oddball paradigm to typically developing children and those with dyslexia (mean age=8.9 years). This study inclined additional tokens with partial ART (40 ms) and FRT (70 ms) durations as ART and FRT deviants. ART deviants did not elicit any mismatch response in either groups. Both partial and full FRT deviants produce mismatch effects at time window 200–300 ms in the control group. In the group of people with dyslexia, mismatch response was elicited only by full FRT deviant. The developmental track of sensitivity was considered in a follow-up study by this research group [56], including three age groups of children (6–8, 8–10, 10–12 years) and adults. Сhildren younger than 10 years showed positive mismatch response between 200 and 400 ms to both ART and FRT conditions. 10–12-year-old children and adults were sensitive only to FRT deviants.

Two papers [57,58] examined the MMN response to the Cantonese syllables with different RTs (/fu/ with RT 120 and 70 ms). In [57] deviant with 70 ms RT elicited more positive mismatch responses (in time windows 150-200 ms after 70 ms RT and 150–238 ms after 120 ms RT) than 120 ms RT deviant. A group of participants with good production and good perception of Cantonese tones showed mismatch responses in both types of counterbalanced contrasts, while a group with good perception but poor production of tones demonstrated mismatch response only when stimulus with a shorter RT (70 ms) was used as a deviant. In the following paper [58] additional group with poor speech production and poor speech perception was also included in the analysis. In this group no significant mismatch responses were observed for both contrasts (120 ms and 70 ms deviants). The amplitude of ERP to stimulus with 70 ms RT at the time window 50–150 ms (P2) was more positive in good production – good perception group than in good perception – poor production group and in poor production – poor perception group. No significant difference between poor production – good perception group and poor production – poor perception group was shown. MMN to 120 RT deviants did not show any significant difference between groups.

Choisdealbha and colleagues [48] provide a longitudinal study of RT perception involving 7 month infants that was followed four months later. Sine tones and speech shaped noise were used as stimuli and RT of standard stimuli was 15 ms and RT of deviants ranging from 161.1 ms to 292.7 ms, in steps of 14.6 ms. It was shown that larger differences in standard and deviant RT produce more positive mismatch response (at time window 300–460 ms). At the same time amplitude of mismatch response becomes more negative in older infants. Stimulus type did not influence the response. The proportion of infants exhibiting MMN or MMR to deviants with different RT was broadly similar across all of the used oddball stimuli.

### ASSR

The study by Van Hirtum and colleagues considered auditory steady-state response (ASSR, an electrophysiological response expressed in neural tuning to the frequency of an auditory stimulus) to RT changes in typical readers and adults with dyslexia [59]. As a stimuli, they used white noise with amplitude modulation at 4, 10, 20, and 40 Hz, with two RT conditions: 10 ms and 30 ms, without affecting the amplitude modulation rate. A sinusoidal envelope modulation was carried out as a baseline condition. As a measure of ASSR strength, the authors considered signal-to-noise (SNR) ratio of ASSR response to an auditory stimulus. ASSR responses were considered in the same EEG frequency ranges as the stimuli.

The study explored auditory processing differences between individuals with and without dyslexia, focusing on their ability to discriminate RTs and perceive speech in noise. Behavioral results revealed that individuals with dyslexia performed similarly to typical readers in discriminating RT and intensity changes and in speech in noise perception. EEG showed distinct neural responses to varying RTs based on the presence of dyslexia. Specifically, participants with dyslexia showed reduced 10 Hz SNRs in 10 and 30 ms RT conditions, lower 20 Hz SNRs for 30 ms RT envelopes, and smaller 40 Hz SNRs for all RTs in the right hemisphere compared to typical readers. No effects were found for 4 Hz SNR. Interestingly, a 30% larger relative increase in RT processing was observed for typical readers in the right hemisphere compared to participants with dyslexia. Increased neural background activity at 40 Hz in the right hemisphere in participants with dyslexia also correlated with poor literacy and phonological skills. In contrast, increased neural background activity of beta activity at 20 Hz was significantly related to better literacy skills. The findings suggest that dyslexia is associated with distinct alterations in neural processing of auditory RTs and increased neural background activity, impacting speech perception and literacy skills. These results are considered within the context of impaired phonological processing in participants with dyslexia, which is manifested in a decrease in SNR by a short RT for stimuli with frequencies of 10 Hz and 20 Hz. The authors also attribute the decrease in SNR in the 40 Hz range to a decrease in phonological processing and the formation of atypical phonological representations. The lack of an effect for 4 Hz is discussed in the context of theta rhythm, which is associated with neural tracking of syllable rhythm, which may not be impaired in participants with dyslexia, unlike phonological processing.

### Rhythmic activity

The study by Le Vere and colleagues considered a mean cortical desynchronization during sleep in relation to different rise times perception [60]. As stimuli, a random noise centered at 125 Hz with two rise time conditions (fast, approximately 3.24 ms, and slow, 7.5 s), were used in the experiment. This study found that both fast-rise and slow-rise auditory stimuli could induce arousal during sleep. However, the effectiveness of the stimulus’s rise time depended on the sleep stage. During non-REM fast-wave sleep, both stimulus types were equally effective. In contrast, during slow-wave sleep, a fast-rise stimulus produced significantly more arousal than a slow-rise stimulus, despite the slow-rise stimulus having a greater total energy and duration. This highlights the importance of rapid onset, rather than overall energy, in capturing attention and causing arousal during deeper stages of sleep.

## Discussion

In the current review, we present the main studies focused on the electrophysiological correlates of RT perception. As can be seen from the given above outcomes, the field under consideration is quite broad and includes a large number of both older classical and modern studies. The main outcomes of the reviewed studies generally are consistent with each other and generalize to the overall framework: The prolongation of the RT leads to a decrease in the amplitude of the main ERP components and an increase in their latencies. Nevertheless, the observed effects may vary and depend on some aspects of the experimental paradigm, that is considered below.

### Stimulation characteristics influencing RT effect

The effect of RT might be influenced by the other stimulation parameters. Below we discuss some of them, such as stimulus presentation rate and stimulus intensity effects on neurophysiological encoding of RT, and suggest optimal stimulation parameters for experimental design. In the following block we also discuss how the stimulus type affects RT decoding as RT might be more crucial for speech sounds. We conclude this section by presenting differential sensitivity to RT changes at different stages of auditory processing.

### Inter-stimulus interval (ISI)

The ISI is an important parameter, especially for ERP studies, due to its effect on the amplitude of the components [61,62]. Different inter-stimulus intervals significantly influence the main ERP components, with component amplitudes generally increasing as the interval lengthens. Moreover, this modulation by stimulus presentation rate varies across developmental stages, reflecting ongoing maturation of neural processing [63]. The reviewed studies used quite variable durations of ISI: from 22 to 457 ms for ABR studies (most often reported as presentation rate); from 500 ms [29] to 14 s [42] for middle and late latency studies; from 10 ms [64] to 1 s [4,54] for MMN/MMR studies. Among ABR and middle and late latency studies included in the review, none compared conditions with different ISI. One MMN study [64] considered the effect of the duration of the interval between stimuli in a pair, which was equal to 10 or 250 ms (design was described in more detail earlier in section 3.9). When the within-pair interval was extremely short (10 ms) RT deviants elicited small MMN and did not elicit any LDN, while in condition with longer interval after previous stimulus both of these components were observed. Another study of this group [35] meanwhile demonstrated the RT effect in a similar short interval: typically developing children showed larger N1 responses to the short RT than to the long RT (10 vs 80 ms). However, in this study, stimuli with different RTs had different frequencies, so the observed results may be related to pitch changes, even authors do not discuss that it could have affected the final outcome. At short intervals after the previous stimulus, masking of the rise time effects may be due to the overlap between the response to the stimulus onset and the response to the previous stimulus offset, or due to insufficient time period for recovery of specific neuronal populations after stimulus-specific adaptation.

In summary of the included studies, we can conclude that an interval of 250 ms is sufficient to detect effects associated with the oddball paradigm (MMN, LDN) [64,65], and an interval of 500 ms is sufficient to observe effects on middle and late latency components[26,29], as at shorter intervals the effect may be masked due to the stimulus-specific adaptation. In ABR studies, the interstimulus interval varied from 22 to 457 ms and did not affect the either amplitude or latency of the component.

### Stimulus type

Studies included in the review used different types of stimuli that can be divided into three main groups: noise bursts (white nose or speech shaped nose), sine tones and speech tokens. Different types of stimuli were compared in the same sample only in two of the reviewed studies [40,48]. In Prasher (1980) study RT effects on N1-P2 amplitude were observed only for tone stimuli, but not for white noise. At the same time, prolongation in N1 latency with RT increasing was observed for both types of stimuli. The author explains this by noting that white noise’s spectral content remains unchanged with varying RT, whereas for tone stimuli shorter RT (e.g., 2.5 ms) can introduce high-frequency transients due to the abrupt onset, making the stimulus sound sharper or harsher. When longer RT were used and tone stimuli were contrasted with speech-shaped noise in the sample of infants, no difference between conditions were observed [48].

A general consideration of the other studies shows that the neurophysiological effects of RT have been successfully detected both in the case of tones and speech stimuli. Some particular discrepancies like in Easwar’s and colleagues’ works[36,39], where significant decrease in N1-P2 amplitude with RT prolongation was observed for syllables but not for 1000 Hz tone burst, are rather due to other stimulation parameters (too tiny difference in RT conditions for the tones). The acceptability of all possible stimulus types for the investigation of RT sensitivity is also supported by a recent longitudinal behavioral study [66]. It was shown that RT sensitivity thresholds measured with different stimulus types are significantly correlated with each other.

The findings from various studies investigating neural responses to speech sound characteristics suggest that formant RT processing often yields particularly distinct or robust effects in neurotypical populations when compared to other groups. For instance, research of Moberly et al., 2014 has indicated that while MMN responses can be elicited by both amplitude and formant RT (ART and FRT) deviants, the brain’s response to formant RT changes is sometimes significantly stronger and more consistently observed across individuals. This pattern of heightened FRT sensitivity in neurotypical group appears to differ from observations in adults with cochlear implants [54], where such a specific enhancement for FRT processing in MMN responses was not evident. Studies involving children also highlight differences based on neurotypical development: typically developing children have been shown to exhibit mismatch responses to a broader range of FRT deviants (including both partial and full changes) compared to children with dyslexia, who may only demonstrate responses to more substantial FRT alterations, implying a more finely tuned FRT processing mechanism in their neurotypical peers [55]. Furthermore, developmental research points to an increasing specialization for FRT in neurotypical individuals. While younger children (e.i., those under 10 years old) might respond to changes in both ART and FRTs, older neurotypical children (e.i., 10–12 years old) and adults tend to show sensitivity primarily to FRT deviants, suggesting a maturation of this specific auditory processing capability [56]. Although not all investigations into RT processing have found group-based differences for every neural component studied, the collective evidence from MMN/MMR studies using specific FRT manipulations, supports the notion that FRT serves as a salient acoustic cue for which neurotypical auditory systems demonstrate more specialized processing.

### Intensity of stimulation

The RT changes accompany the total changes of the stimulus energy, such as intensity and also significantly influences perceived loudness of a sound. This effect is associated with the reduction of the plateau duration (period at which the stimulus has maximal intensity) if RT is increased. Therefore, stimuli with the same duration but different RTs will have different cumulative intensity and can be perceived as stimuli of different loudness. In order to distinguish RT and intensity effects, Thomson and colleagues (2008) used two types of deviant stimuli: deviant with a shorter RT (15 ms) than in standard stimulus (50 ms) and deviant with the same short RT but matched with the standard stimulus in intensity. When intensity of RT deviant was matched, no significant mismatch response was found, while traditionally used RT deviant, that is unmatched in intensity, produced some MMN (but it was not significant). However, in the opposite condition, when the standard and deviant stimuli were switched, matched in intensity deviant with longer than standard RT (50 ms) elicited MMN. Thus, MMN is sensitive to the RT per se, but dependent on the context. In ERP study by Kodera and colleagues (1979), stimuli with different RT were matched in intensities. Despite this, the authors obtained significant RT effects for latency and amplitude of ABR and main middle and late latency components (except N1-P2 peak-to-peak amplitude). The absence of an effect for the N1-P2 component is quite unusual, as most studies have associated this component with the RT effects. This effect may be related to the fact that when intensity is equated, the main effects of RT are related to the signal detection time. When stimuli are equalized in intensity, the point at which stimuli with different rise times begin to be perceived as equivalent occurs earlier. In this case, such difference in stimulus detection time could be less critical for effects on later components, especially when the difference between the RT conditions is small (15 ms in Kodera work). In relation to ABR, due to the even smaller difference in the durations of the RTs studied, this problem is relevant only in the sense that some RT+intensity combinations are below the response elicitation threshold and, accordingly, cannot be compared.

RT discrimination thresholds are influenced by the overall stimulus intensity [67]. While many studies in the literature, particularly those focusing on language development or clinical applications, control for intensity by presenting sounds at comfortable listening levels, the general psychophysical principles suggest an intensity dependence. It is generally understood that at very low sound levels, the auditory system has fewer activation, leading to poorer discrimination. As intensity increases, more auditory nerve fibers are recruited, providing a more robust and detailed representation of the sound’s temporal envelope, which can lead to improved RT discrimination. This complex interaction between intensity and temporal processing is crucial for understanding how we perceive sounds in various real-world environments. So, it is important to consider whether intensity is an essential characteristic for detecting RT effects on the ERP. Some classical auditory perception studies have investigated the effects of RT at different intensities. Onishi and Davis (1968) have found that the prolongations in N1 latency and decrease in N1-P2 peak-to-peak amplitude with RT increase were observed in all intensity levels (45, 65, and 85 dB), notable that latency increase was particularly marked at the lowest (45 dB) intensity level. However, in the second part of their study, an intensity condition of 15 was used, at this condition response was above noise level for only one participant for the rise time of 3 ms, and only for two participants for rise time of 30 ms. Nevertheless, the authors report that the amplitude of N1P2 was systematically lower at 3 ms RT than at 30 ms RT, at all intensity levels. No interaction between RT and intensity effects was found in the Skinner and Jones study [30]. However, it should be noted that no statistical analyses were performed in the two given studies. In Prasher study (1980) the observed RT effects did not differ between intensity conditions (80 and 40 dB SPL). Overall, no clear association between the observed RT effects and intensity was reported in the included in the review studies. Thus, modulation of amplitude and latency by the RT duration can be detected even at low intensities (15–45 dB SPL), especially if the difference between the RT durations is large enough [27,30,37,42].

Thus, the findings of this systematic review suggest that variations in intensity across different RT conditions did not influence the neurophysiological discrimination, but could potentially confound the interpretation of results. However, there is no effective way to separate the effect of RT and cumulative intensity as inevitable matching of cumulative intensity of stimuli with different RT lead to their difference in maximal intensity.

### Differential sensitivity to the rise time duration changes across stages of auditory processing

The reviewed studies cover a very wide difference in RT from microseconds to hundreds of milliseconds, all of them indexing some aspects of the natural environment. Fast abrupt changes activate the alerting system and also important for consonant discrimination, RT between 15 to 100 ms are crucial for vowel perception, while RT changes in slowed rate represent the background auditory scene analysis, emotional component of speech and prosody [68–70].

A summary of differences in RT durations for different ERPs can be seen on Figure 3. In most ABR-studies RT differences varied from tens of microseconds (the smallest value 90 µs or 0.09 ms in Salt & Thornton study) to 7 ms, while in some studies this difference reached 15 ms. Noteworthy, for each contrast the significant changes in ABR latencies were found. Middle latency components were studied only in two studies with the RT range from 0.49 ms to 24.99 ms and the components modulation were demonstrated for all used contrasts. The range of RT differences, which showed significant results for late latency (components was from 4.99 ms to 500 ms, in most cases considered as N1P2 peak-to-peak amplitude). The optimal RT ranges also differed within late latency components being smaller for N1, and larger for P2 (>30 ms). RT differences less than 4.99 ms (namely 2.49 ms in Prasher study, 1980) and more than 500 ms (in Elfner study, 1976) showed no significant changes in amplitude or latency.

**Fig. 3.**
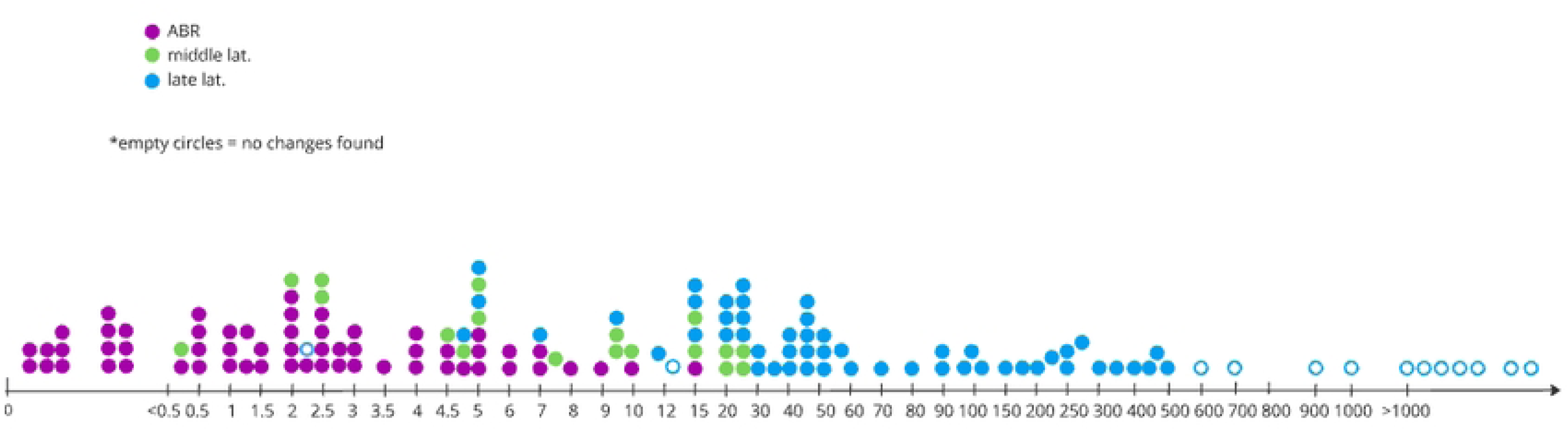
Differences in rise time durations (in ms) in studies observed.

In most recent studies on the RT perception, a RT of 15 ms is usually used as the minimum value [29,33,47,48,65]. This concerns those evoked potentials that arise in the cerebral cortex as opposed to those that are associated with activity, for example, in the brainstem, namely, the ABRs, which are produced on the earlier stages of auditory processing and are not accessible for behavioral discrimination. In auditory discrimination tasks, 15 ms is widely utilized as a standard reference point which is compared to a longer RT. This standardization allows for consistent experimental protocols and comparable results across studies. According to research on auditory processing and phonological development, this parameter provides an optimal baseline for discrimination tasks. In RT discrimination tasks, researchers typically present target sounds gradually logarithmically ranging from 15 ms (the standard) to longer durations. This methodological approach has been validated through multiple studies, which explore relationships between RT sensitivity and reading development [29,48,71]. Research has also demonstrated that 15 ms RT effectively balances the need to avoid spectral splatter while maintaining temporal precision in auditory stimuli. This technical consideration is crucial for creating clean experimental stimuli without introducing confounding variables [72]. The 15 ms RT parameter aligns with critical amplitude RTs in the speech envelope that significantly impact speech intelligibility. This connection to natural speech processing makes it particularly relevant for studying both typical and atypical language development, as demonstrated in research on auditory temporal processing and phonological awareness [73].

Thus, the optimal time window for detecting RT effects varies depending on the level of processing. For brainstem structures, the response of which is detected by ABRs, it is very short. For subcortical structures it is longer, and for cortical structures (late potentials) it is the longest and in most works is not less than 15 ms. While ABRs are remarkably sensitive to RT differences as tens of microseconds, reflecting their role in processing the immediate onset of sound, subcortical structures demonstrate a longer optimal window. This extended window is evidenced by studies on middle latency components, which showed modulations across a range of RT differences from 0.49 ms to 24.99 ms. This suggests that subcortical processing integrates auditory information over a slightly more prolonged period than the brainstem, allowing for the detection of RT effects that unfold over hundreds of microseconds to several milliseconds. Further extending this temporal integration, cortical structures, responsible for late potentials, exhibit the longest optimal window for RT effects. In the majority of research, this window is not less than 15 ms. This is because late cortical components are involved in higher-level processing, where the brain is analyzing more complex features of the sound stimulus. Consequently, to observe significant effects, a more substantial difference in the RT between conditions – typically exceeding 15 ms – is required for the distinctions to be sufficiently pronounced to modulate cortical responses. This extended temporal requirement reflects the cortical areas’ role in integrating information over longer durations for perceptual discrimination, language processing, and cognitive functions, where the precise, rapid onset information critical for brainstem responses has already been processed and is being interpreted within a broader temporal context.

### Developmental changes in electrophysiological markers of rise time perception

The main ERP components undergo significant developmental changes across the lifespan, reflecting the maturation of the auditory processing system. Already the earliest, subcortical auditory responses like ABR demonstrate significant age-related changes, with research showing progressive increases in peak latencies of waves I, III, and V through ontogenesis [74]. A systematic search, however, found only one RT study in children that examined ABR: the results for infants in this study were no different from those for adults [14]. No RT studies for children concerning middle latency potentials were found.

Late latency components like N1 demonstrate prolonged development. In early childhood, auditory evoked potentials are dominated by the P1 and N2 components, while the N1 and P2 components response becomes more prominent with maturation at about 10-12 years of age. Also latency of such components decreased with age. The MMN component, which reflects automatic auditory change detection, develops from positive to negative through childhood. MMN has been directly linked to auditory discrimination abilities that evolve throughout the lifespan [75]. These developmental changes in ERPs considered in relation to RT perception have important implications for understanding both typical auditory development and disorders characterized by impaired rise time processing.

Studies of neurotypical adults demonstrated that the main effects of RT are observed on N1 and P2 components. However, in young children these components are not yet fully developed [76], which suggests that the typical pattern of ERP components modulation by the rise time duration may change with age. Only three studies included in the review examined the RT effects of main ERP components in children [29,35,41]. Two of them considered RT effects on N1–P2 peak-to-peak amplitude in children samples [35,41]. In the study by Carpenter & Shanin (2013) [41] decreasing N1-P2 amplitude was observed only after 6 years, while 4–5 years old children did not demonstrate any RT effects, which may be related to the poor development of this component in this age group. In Hämäläinen (2007) [35] study, typically reading children (aged 8.8–10.5 years) demonstrate N1 decrease with RT prolongation. The third of the included studies concerning RT in children [29] showed that in children the RT related effects can be transferred to adjacent components (P1 and N2), which probably absorb some functions from not fully developed N1 and P2 components. Thus, in children (aged 7–11 years), P1 demonstrates a latency delay with RT increasing. It was also shown that at these ages RT effects are observed for the N2 component, which demonstrates a decrease in amplitude and prolongation of latency with RT increasing.

Age-specific phenomenon was also observed for studies carried out in the oddball paradigm. Generally, the latency of the difference-wave mismatched response decreases with age. Decreasing in MMN latency in older children is a common fact associated with age-related reduction in latency of ERP components [77]. This is also typical for the mismatch response evoked by a deviant stimulus with a different RT, which can be seen in studies, included in this review. Six of the oddball studies included in this review also examined children. In particular, in studies including adult participants, MMN to RT changes was observed in a time window from 150 to 350 ms [4,52,54,57,58]. In children, this time window can be shifted and is considered as a mismatch response [47,56,56,65]. At 11 years the peak of mismatch response appears at 256 ms [65], while at age of 3 years – at 346 ms [47]. In infants, the response time window is shifted even more: from 300 ms to 460 ms [48]. Also, age could affect MMN amplitude, which tends to be larger in children than in adults [77,78]. From the works presented in the review we can also observe the change of configuration of the mismatch response on difference wave, which becomes more negative in older participants. The beginnings of these changes can already be observed in infants, but the response remains positive (Choisdealbha et al., 2022). At around 10 years, the positive MMR is replaced by a negative response, known as MMN [56].

Behavioral studies have established that discriminatory sensitivity to acoustic RT changes throughout ontogenesis [71]. In particular, six-year-old children have significantly lower RT thresholds (83.6 ms for sine tones, 150.26 ms for noise, 61.06 ms for speech token) than four-year-olds (205.37 ms for sine tones, 257.69 ms for noise, 105.91 ms for speech token). However, the neurophysiological underpinnings of this developmental trajectory remain insufficiently characterized, largely due to a scarcity of research focused specifically on pediatric populations. The existing literature provides glimpses into these developmental shifts. For example, Peter et al. (2018) studied speech tokens and demonstrated a clear age-dependent pattern where children younger than 10 showed a positive mismatch response to both ART and FRT deviants, whereas children aged 10-12 and adults exhibited sensitivity primarily to FRT changes. This suggests a maturational process that refines neural specialization for speech sounds. Consequently, while it is clear that neural sensitivity to RT evolves with age, a comprehensive map of how specific neurophysiological markers for different types of RT develop in children is still needed.

Thus, it is important to take into account that the main target components of ERPs showing RT effects quite significantly varies with age. Thus, the ERP configuration in each age group has to be considered in order to more accurately interpret the RT effects and to be able to compare the results of different studies.

### Rise time perception and speech disorders

Eight of included studies were focused on the neurophysiological specifics of RT processing in speech disorders (particularly dyslexia). Four of them consider MMN or MMR measured in oddball paradigm as possible neuromarker of RT processing in speech impediment groups [47,55,64,65], three works consider late latency ERP components (such as P1, N1, P2 and N2 [29,34,35] and one work was focused on ASSR [59]. The observed results were very controversial. Several studies indicated that some patterns of ERP modulation by RT that were observed in the typically-developing group were not present in groups with reading disabilities [35,47,55]. In contrast, other studies have shown that RT-related changes that were not characteristic of the typically-developing group are observed in reading disorders [29,34,64]. Alternatively, there may be no differences between groups at all [65].

Research into the RT processing of individuals with dyslexia, particularly concerning the MMN and other ERP components in response to changes in stimulus RT, has yielded varied results. One key finding from Hämäläinen and colleagues (2008) indicated a larger MMN to RT changes in children with reading difficulties when preceding stimuli were sufficiently separated. However, this observation was tempered by the suggestion that an elevated N1 component in dyslexia [79] might contribute to this effect through overlap. In subsequent work, Hämäläinen et al. (2011) noted that individuals with dyslexia exhibited less negative amplitudes of the T-complex with prolonged RTs, which they attributed to an enlarged Tb component, potentially enabling better detection of RT effects despite other processing differences. Other studies also explored this area, with Stefanics (2011) reporting the increased P1 latency and bigger fronto-temporal N1c amplitude (for 90 ms RT in dyslexia group vs 15 ms RT in TD group) for RT changes in children with dyslexia as compared to neurotypical controls.

Despite these findings, the Peter (2016) study introduced a nuanced perspective, suggesting that the perception of formant RT differences might be a more informative marker for dyslexia than amplitude RT differences. This raises questions about the overall utility of the MMN response to amplitude RT deviants as a definitive indicator of dyslexia. Further insights come from Plakas (2013), whose study on three-year-old children revealed a critical distinction: MMR in response to a 90 ms RT tone within a stream of 15 ms RT tones was exclusively observed in typically-developing children without a familial risk of dyslexia. Children with a familial risk, regardless of a dyslexia diagnosis, did not exhibit this response. This result points towards a genetic predisposition to dyslexia, rather than merely reflecting the diagnosis itself, and suggests that these early childhood MMR differences might signify a general susceptibility to phonological challenges that could be mitigated by the development of other language skills in older age. Collectively, these studies underscore the complex and multifaceted nature of auditory processing deficits in dyslexia, highlighting the need for continued research to refine our understanding and identify robust diagnostic markers.

### Directions for further research

As previously mentioned, there are certain areas in RT research where the findings are largely consistent, such as decrease in amplitude and increase in latency for the main components of ABRs and ERPs. Conversely, the existing results do not permit us to draw clear conclusions about the mechanisms underlying RT effects, particularly in studies related to MMN, LDN, and research involving individuals with dyslexia. It is worth noting that many of the studies yielding consistent results were conducted quite some time ago, which suggests that new methodological approaches and more advanced technology could potentially reinterpret these findings. This is particularly relevant for ABR-studies. Due to methodological limitations, including outdated technology, small sample sizes, and manual measurements, existing ABR rise time studies, the most recent of which dates back to 1996 with a significant portion conducted in the 1970s-1980s, may be biased, highlighting the need for replication with modern equipment, larger samples, and design-appropriate statistics. Another clear gap in knowledge that is evident at Fig.2, is the absence of research on the main ERP components after 2014, with the field dominated by an oddball paradigm. While oddball paradigm allows assessing the neurophysiological response to the sound differences, it has several drawbacks, such as increased duration of stimulation, low signal to noise ratio, ambiguity of results interpretation and so on [49,80]. Thus, the recent neurophysiological studies on RT can be largely supplemented by the analysis of main ERP components in response to repetitive stimulation.

In this review, we aimed to observe all RT electrophysiological human studies. However, studies in the children population, especially for basic ERP components, such as P1, N1, P2 and N2 are lacking, while investigation of the developmental trajectory for the rise time neurophysiological sensitivity is crucial for linking it with speech perception ability. A similar gap is seen for the studies in the clinical group, only participants with speech and reading difficulties were found to be included in the review. At the same time, impaired ability to process phonological features of auditory stimuli is observed not only in patients with other speech and language disorders, but also in patients with developmental disorders, primarily with ASD [81]. In particular, it was reported about an auditory temporal-envelope resolution deficit in this population [82], which might cause the delay of phonological categories development in children with ASD [83]. Temporal resolution of the auditory cortex is often assessed using amplitude modulated stimuli [84], and such a characteristic as RT can be also differently processed in people with ASD. Thus, investigation of RT neurophysiological discrimination in people with ASD seems like a logical continuation of such research in clinical groups.

### Limitations

Several limitations of this review should be acknowledged. First, our search strategy was restricted to publications in English and to major electronic databases, which may have introduced language and publication biases, potentially omitting relevant studies published elsewhere.

Second, there was substantial methodological and conceptual heterogeneity across studies in some sections. This heterogeneity precluded a quantitative meta-analysis and necessitated a narrative synthesis, which is more susceptible to subjective interpretation. To enhance objectivity, we used a pre-defined framework for synthesis and independent assessment by two reviewers.

Third, the methodological quality of the primary studies was sometimes suboptimal, with frequent unclear or high risk of bias, as assessed by the OHAT tool. Consequently, the overall confidence in the body of evidence for outcomes in some sections, as per our OHAT-based assessment, was not high enough. This fundamentally qualifies the strength of our conclusions, which should be viewed as hypothesis-generating rather than definitive.

Finally, despite our comprehensive search, the possibility of unpublished null results (publication bias) remains a concern that could skew the presented narrative towards an overestimation of reported effects.

## Author contributions

**V.M.**: Conceptualization, Full-text screening, Analysis, Writing – original draft, Writing – review and editing, Visualization. **D.K.**: Conceptualization, Abstract screening, Full-text screening, Analysis, Writing – original draft, Writing – review and editing. **A.R.**: Abstract screening, Full-text screening, Writing – original draft, Writing – review and editing. **O.S.**: Conceptualization, Writing – review and editing, Supervision

